# Synthetic mammalian signaling circuits for robust cell population control

**DOI:** 10.1101/2020.09.02.278564

**Authors:** Yitong Ma, Mark W. Budde, Michaëlle N. Mayalu, Junqin Zhu, Andrew C. Lu, Richard M. Murray, Michael B. Elowitz

**Affiliations:** Division of Biology and Biological Engineering, California Institute of Technology, Pasadena, CA 91125, USA; Division of Engineering and Applied Science, California Institute of Technology, Pasadena, CA 91125, USA; Department of Biology, Stanford University, Stanford, CA 94305, USA; Howard Hughes Medical Institute, California Institute of Technology, Pasadena, CA 91125

## Abstract

In multicellular organisms, cells actively sense and control their own population density. Synthetic mammalian quorum sensing circuits could provide insight into principles of population control and extend cell therapies. However, a key challenge is reducing their inherent sensitivity to “cheater” mutations that evade control. Here, we repurposed the plant hormone auxin to enable orthogonal mammalian cell-cell communication and quorum sensing. We designed a paradoxical population control circuit, termed *Paradaux,* in which auxin stimulates and inhibits net cell growth at different concentrations. This circuit limited population size over extended timescales, of up to 42 days of continuous culture. By contrast, when operating in a non-paradoxical regime, the same cells limited population growth, but were more susceptible to mutational escape. These results establish auxin as a versatile “private” communication system, and demonstrate that paradoxical circuit architectures can provide robust population control.

## Introduction

Cells use intercellular communication systems to sense and control their own cell population density. In microbial communities, cells secrete diffusive signals to coordinate cooperative behaviors through the process of quorum sensing (Waters and Bassler, 2005; Papenfort and Bassler, 2016). In multicellular organisms, intercellular communication is essential to enable precise developmental patterning (Gibb, Maroto and Dale, 2010; Bier and De Robertis, 2015), control immunological responses (Muldoon *et al*., no date; Boyman and Sprent, 2012; Chen *et al*., 2015), and coordinate organism-level physiology (Keener and Sneyd, 2009, chap. 16). Synthetic intercellular communication systems could similarly allow the engineering of inherently multicellular behaviors not possible with cell-autonomous circuits (Toda *et al*., 2018). For example, in bacteria, foundational synthetic biology studies showed how coupling quorum sensing systems to cell death could enable bacteria to limit their own population size (You *et al*., 2004; Scott and Hasty, 2016), or drive synchronized oscillations of drug release for therapeutic applications (You *et al*., 2004; Scott and Hasty, 2016).

In mammalian cells, an orthogonal, or “private” communication channel that allows specific communication between cells could enable engineering of analogous circuits (Figure 1A). Mammalian cells have been engineered to produce, sense, and process signals from natural ligands by rewiring signaling pathways such as Nodal-Lefty, Sonic Hedgehog, and Notch (Matsuda *et al*., 2012; Sekine, Shibata and Ebisuya, 2018; Li and Elowitz, 2019), and by repurposing the amino acid tryptophan as a signaling molecule (Bacchus *et al*., 2012). However, these approaches are not orthogonal to endogenous systems. On the other hand, synNotch receptors allow multiple orthogonal communication channels, but depend on cell contact interactions to mediate diffusible signaling (Toda *et al*., 2020).

**Figure 1:**
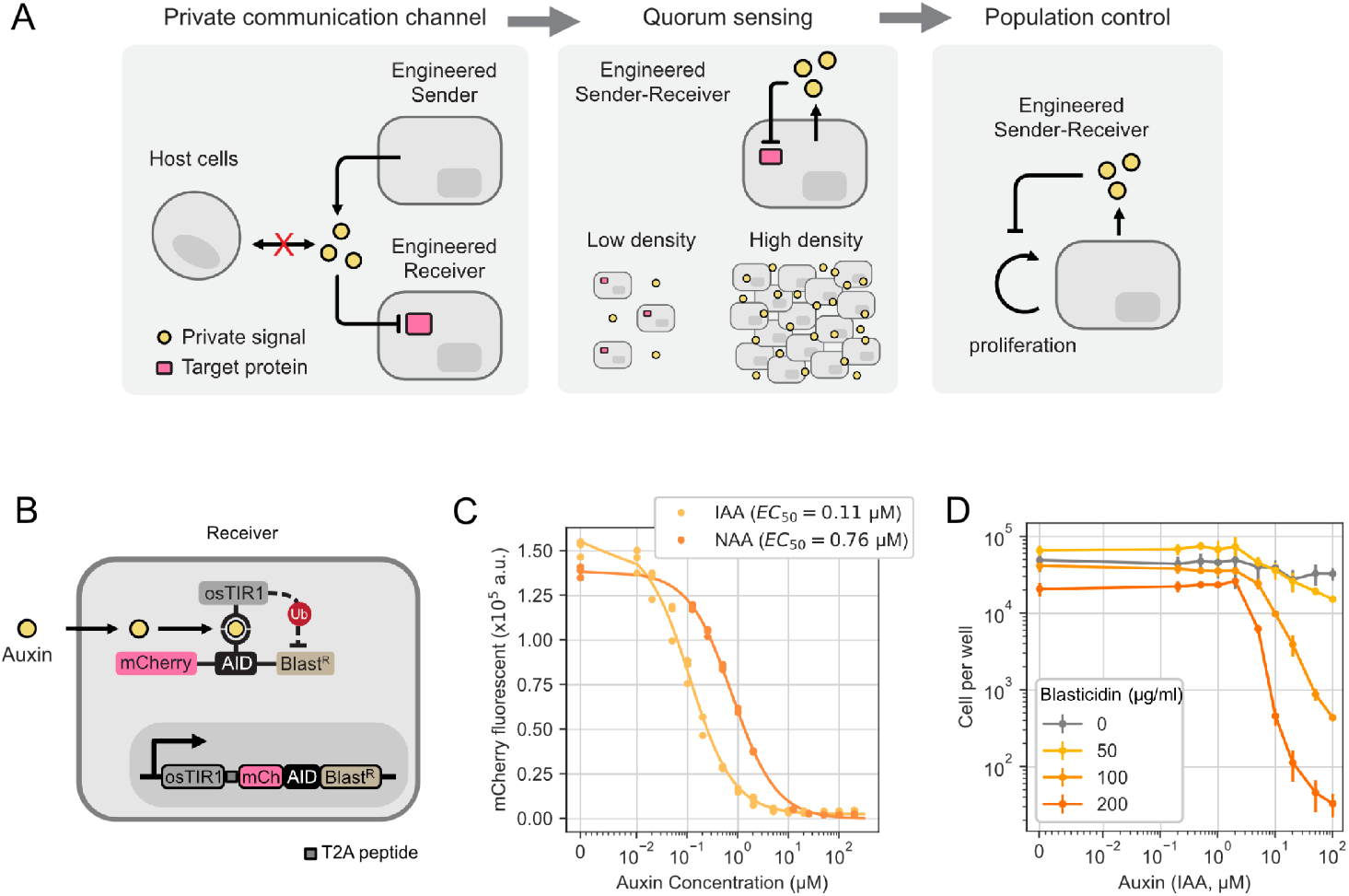
The plant hormone auxin allows private-channel mammalian cell-cell communication. (A) Left: an ideal mammalian private-channel communication system would allow engineered cells to send and respond to an orthogonal signal that does not interact with host cells. Middle: Engineered cells that can send and receive the signal simultaneously can respond to their own population size. Right: coupling the sending-receiving function with cell survival in a negative feedback loop enables population control. (B) Auxin Receiver cells constitutively express a fluorescent target fusion protein, mCherry-AID-BlastR, as well as the F-box protein osTIR1. In the presence of auxin (yellow circle), osTIR1 and the AID-tagged target protein assemble into an SCF complex, which allows ubiquitylation of the target protein, leading to target degradation. Both proteins are encoded on a single transcript, with an intervening T2A ribosomal skip sequence (grey square) to yield separate proteins (Szymczak *et al*., 2004). (C) Auxin regulates intracellular protein levels. The response of mCherry fluorescence in Receiver cells (B) to two different species of auxin (IAA and NAA) was measured after two days of treatment (dots). The responses follow Michaelis-Menten kinetics (fitted lines, Method), with indicated EC_50_ values. (D) Auxin regulates cell density. Cells were treated with a combination of IAA and blasticidin at different concentrations for four days and passaged once. Cells were counted by flow cytometry (n=3, error bar = standard deviation). In (C) and (D), the x axis uses a symmetric log (symlog) scale to include a value of 0.

The ideal private communication system for mammalian population control would use a diffusible signal, avoid undesired interactions with non-engineered cells, permit external control over the strength of signaling, and operate in a broad variety of cell types. It should also allow direct and rapid control of diverse target protein activities to allow flexible interfacing within cells, and exhibit minimal immunogenicity to facilitate potential biomedical applications. Auxins, a class of plant-specific hormones that coordinate growth and behavior including root initiation, embryogenesis, and tropism (Tanaka *et al*., 2006), represent an excellent candidate for this role. Molecularly, auxin induces protein-protein interactions between the F-box transport inhibitor response 1 (TIR1) protein and its target proteins. This leads to the assembly of a Skp, Cullin, F-box containing (SCF) complex, which in turn recruits E2 ubiquitin ligases that target specific proteins for degradation (Reitsma *et al*., 2017). Because TIR1 and its targets are absent in mammals, auxin does not regulate endogenous mammalian proteins. However, ectopic expression of TIR1 from rice (osTIR1) is sufficient to confer auxin-dependent degradation of proteins engineered to contain a minimal auxin inducible degron (mAID, or AID for simplicity in this paper) (Nishimura *et al*., 2009; Natsume *et al*., 2016). Thus, auxin is orthogonal to endogenous mammalian pathways, but can enable direct control of cellular activities through engineered protein targets. Additionally, in yeast, ectopic expression of bacterial indole-3-acetic acid hydrolase was shown to catalyze auxin production from an inactive precursor indole-3-acetamide (IAM), allowing control over auxin production (Khakhar *et al*., 2016). Nevertheless, a full auxin sending and receiving signaling system, which is necessary for population control, has not been established in mammalian cells.

A critical challenge for any population control circuit is evolutionary robustness. By limiting growth, a population control circuit inherently selects for “cheater” mutations that escape regulation. In bacteria, toxin-antitoxin systems and periodic strain replacement can prevent cheater escape (Stirling *et al*., 2018; Liao *et al*., 2019). However, these systems use components that do not function in mammalian cells or are not cell-autonomous. In the mammalian context, a seminal analysis of natural cell population size control systems in cytokine and glucose sensing circuits revealed a paradoxical architecture, in which a single signal stimulates both proliferation and death of the same target cell population, to actively select against cheaters (Hart *et al*., 2014). In this paradoxical design, mutations that diminish signal sensing lead to cell death and are eliminated (Karin and Alon, 2017). Despite its power and elegance, the paradoxical architecture has not, to our knowledge, been demonstrated synthetically in living cells.

Here, we engineer the auxin pathway to act as a private mammalian communication channel, and use it to construct and analyze synthetic population control circuits with different architectures. Combining auxin-synthesizing enzymes and auxin transporters, and employing alternative auxin precursors, we show that the auxin pathway can be used for synthetic quorum sensing in mammalian cells. Using this pathway, we constructed and compared negative feedback and paradoxical control systems that regulate their own population size through auxin quorum sensing. While both circuits limit population size at early times, the paradoxical system enhances evolutionary stability, as predicted theoretically, extending the duration of population control. Together, these results provide a versatile, diffusible synthetic signaling module for private-channel communication and demonstrate how paradoxical control schemes can enhance the evolutionary stability of population control systems.

## Results

### Engineered mammalian cell lines sense, respond to, and produce auxin

To establish and characterize auxin regulation of mammalian cell growth, we coupled auxin sensing to drug resistance and fluorescence. Specifically, we fused blasticidin S deaminase (BlastR) (Kimura, Takatsuki and Yamaguchi, 1994), whose protein product is necessary for survival in the presence of blasticidin, to AID and mCherry domains, allowing auxin-dependent degradation and fluorescent readout of protein concentration, respectively. We then stably integrated this chimeric gene, along with a constitutively co-expressed osTIR1, in CHO-K1 cells to create an auxin-sensitive “Receiver” cell line (Figure 1B).

To validate auxin regulation of mCherry-AID-BlastR, we cultured Receivers in media containing different concentrations of two auxin variants: either the major natural auxin, indole-3-acetic acid (IAA), or a synthetic auxin analog, 1-napthalenatic acid (NAA) (Figure 1C). Both auxins reduced mCherry fluorescence in a dose-dependent manner, with EC_50_ values of 0.11 μM and 0.76 μM, respectively. Addition of IAA to media containing blasticidin was sufficient to degrade BlastR and inhibit cell survival (Figure 1D). This effect was dose-dependent with both blasticidin and IAA. Comparing fluorescence of mCherry-AID-BlastR in Figure 1C with cell survival in Figure 1D shows that a small amount of mCherry-AID-BlastR is sufficient to enable survival in blasticidin. These results confirmed that the AID domain and osTIR1 together are sufficient to enable net growth regulation by auxin (IAA) in mammalian cells.

In addition to sensing, a population control system requires that cells produce auxin at levels sufficient to trigger responses in receiving cells (Khakhar *et al*., 2016). Auxin can be synthesized in two enzymatic steps: (1) oxidation of L-tryptophan to indole-3-acetamide (IAM) and (2) hydrolysis of IAM to IAA (Figure S1A) (Sitbon *et al*., 1992). By itself, the second enzymatic step can produce either IAA or NAA from precursors IAM or 1-naphthaleneacetamide (NAM), respectively, enabling direct control of auxin production (Figure 2A) (Kawaguchi *et al*., 1991). To identify enzymes that efficiently catalyze this reaction, we compared thirteen indole-3-acetamide hydrolases from bacteria and plants (Figure S1B, left), transiently expressed them individually in Receivers, and measured their ability to downregulate AID-tagged mCherry fluorescence by flow cytometry.

**Figure 2:**
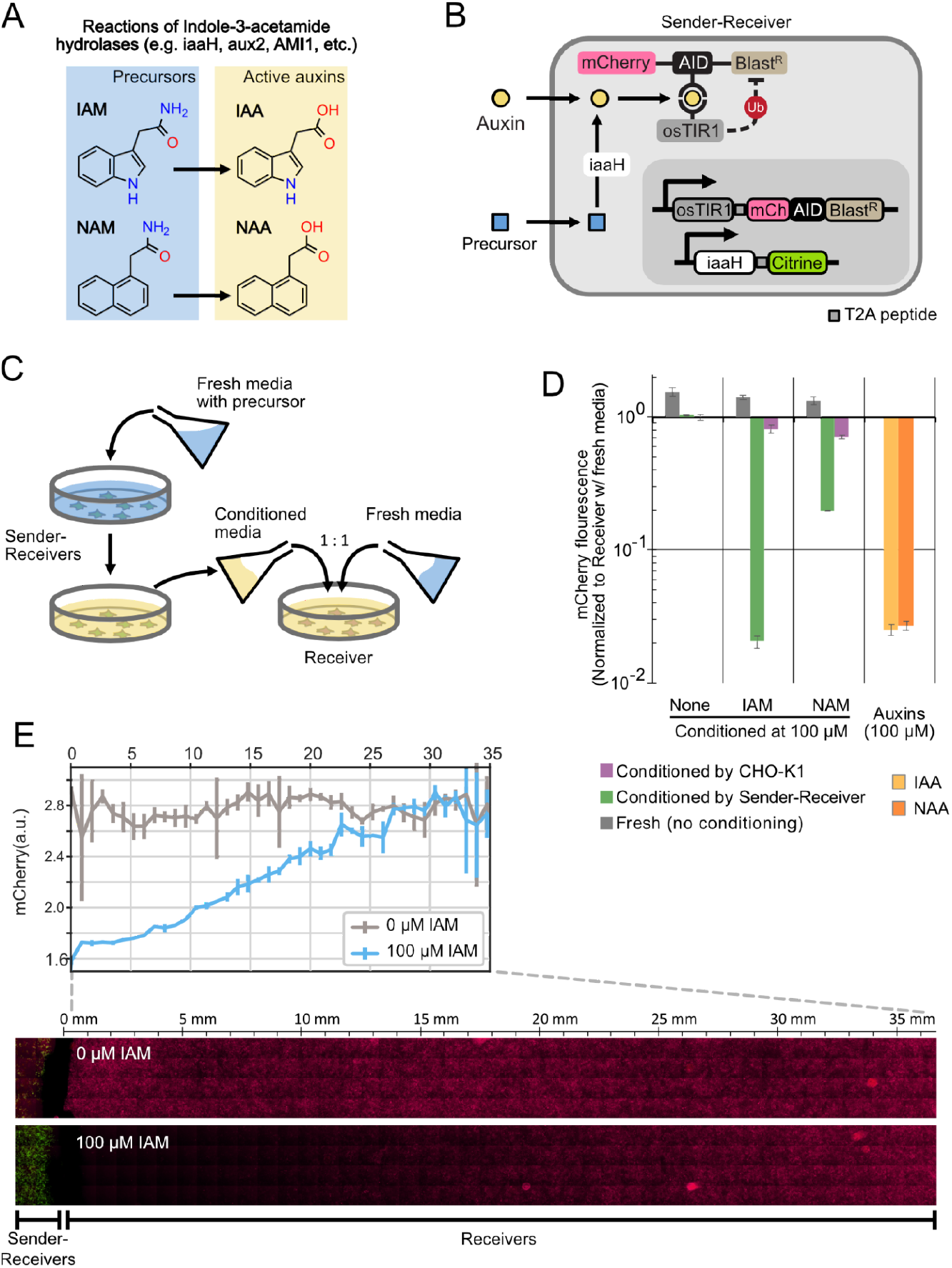
Sender-Receiver cells produce and respond to auxin. (A) Indole-3-acetic acid hydrolases such as iaaH, aux2, and AMI1 hydrolyze inactive auxin precursors (IAM and NAM) to their respective active form (IAA and NAA). (B) Stable expression of iaaH in Receiver cells allows them to produce auxin from precursors (blue square). (C) Conditioned media experiment (schematic). Fresh culture media with or without precursors was added to plated sender cells, collected, mixed at 1:1 ratio with standard fresh media, and then applied to receiver cells. (D) iaaH can produce IAA and NAA auxins from IAM and NAM precursors, respectively. Media with or without precursors were conditioned by Sender-Receivers or standard CHO-K1 cells for 48 hours and applied to Receiver cells for another two days. Receivers cultured with fresh media with or without auxins were also assayed as controls. Data are normalized to Receiver cell fluorescence treated with media conditioned by CHO-K1 cells. Error bars represent standard deviation of 3 replicates. (E) Auxin senders can generate an auxin gradient. Sender-Receivers (green) were seeded within a 7mm x 7mm square at the edge of a 60mm dish, and Receivers were plated everywhere else (Receiver region). One day after plating, the media was replaced with fresh media containing low-melting-point agarose, with or without IAM (Methods). Plates were imaged after two additional days of culture. Inset on top: Quantification of the average pixel intensity of mCherry expression in cells shows that mCherry inhibition depends on distance from Sender-Receiver region. Error bars denote standard deviation of the four images making up each column in the mosaic.

Three enzymes reduced mCherry to levels comparable to that produced by addition of IAA itself (Figure S1B). Among these, we selected *A. tumefaciens* iaaH for further use. We stably integrated it in Receivers to create a Sender-Receiver cell line (Figure 2B). After 2 days of culturing Sender-Receivers in media containing the IAM precursor, the resulting conditioned media, diluted into an equal volume of fresh media for optimum cell growth, reduced mCherry-AID-BlastR to levels comparable to those generated by saturating concentrations of IAA (Figures 2C and 2D). The Sender-Receiver line was also able to produce the auxin NAA (which, as shown below, has some advantages compared to IAA) from its corresponding precursor NAM (Figures 2D), albeit with a diminished response compared to NAA, consistent with the higher EC_50_ of NAA compared to IAA. These results show that iaaH expression in mammalian cells can efficiently produce both auxins from corresponding precursors.

The ability to synthesize auxin in mammalian cells without exogenous precursors could facilitate in vivo applications of population control circuits. Bacterial auxin (IAA) synthesis pathways use a tryptophan 2-monooxygenase, iaaM (also known as aux1 or TMO), to synthesize IAM from L-tryptophan (Figure S1A). Sender-Receivers expressing iaaM (from *P. savastanoi*), cultured in media without precursors, produced auxin concentrations in conditioned media that were sufficient to degrade the auxin reporter in Receiver cells (Figure S1C). Furthermore, iaaM-expressing Receivers produced precursor in conditioned media, allowing auxin production by Sender-Receivers (Figure S1C). These results demonstrate the two step iaaM-iaaH auxin synthesis pathway can operate in mammalian cells without exogenous precursors. However, in the following experiments, we used iaaH with added precursors to allow external control of auxin synthesis.

Population control requires integrating cell density over an extended spatial scale. To estimate the spatial range of auxin signaling, we seeded a field of Sender-Receivers adjacent to a larger region of Receivers, and applied media in a 1.5% agarose gel to prevent non-diffusive transport (Methods). After 48 hours, mCherry fluorescence was reduced in Receivers proximal to the Sender-Receiver region, forming a long-range gradient. This effect occurred in the presence, but not the absence, of the IAM precursor, consistent with a dependence on IAA production (Figure 2E, bottom). Image analysis revealed auxin response in Receivers declining to half its maximum value at a length scale of 15.6±0.85 millimeters (Methods), or approximately 750 cell diameters, from the source region, within 48 hours (Figure 2E, top), consistent with expectations for a small molecule the size of auxin diffusing in buffered solutions (Robinson, Anderson and Lin, 1990) (Figure S1D and STAR Methods). These results show that auxin can provide information about global cell density over an extended region, and further demonstrate the possibility of using auxin as a synthetic gradient-forming morphogen for applications in synthetic developmental biology (Teague, Guye and Weiss, 2016).

### Engineered cell lines sense population density

We next asked whether Sender-Receiver cells could sense their own population density by producing auxin at a rate proportional to population size and sensing auxin concentration in the local environment (quorum sensing, Figure 1A, middle panel). We analyzed the dependence of reporter expression on cell population density in the presence of concentrations of each precursor (NAM and IAM) sufficient to generate saturating levels of auxin. With the NAM precursor, reporter fluorescence decreased in response to increasing population density (Figure 3A, darker blue line). By contrast, addition of IAM generated a strong, but density-independent, decrease in fluorescence (Figure 3A, light blue line). We reasoned that this density-independence could result from IAA’s limited membrane permeability at neutral pH (Raven, 1975), potentially causing newly synthesized IAA to accumulate intracellularly. Though the rate of exchange is sufficient to fully induce receiver cells (on the timescale of days, Figure 2D and 2E), it could be insufficient to prevent intracellular accumulation of IAA due to the faster rate of auxin production by iaaH (typically several μM per minute) in the same cell (Mishra *et al*., 2016). To overcome this issue, we stably expressed the auxin exporter PIN2 from *Arabidopsis thaliana* (Petrásek *et al*., 2006) in Sender-Receiver cells (Sender-Receiver-PIN2 cells) (Figure 3B, right). PIN2 expression produced a modest decrease in auxin sensing, suggesting the transporter was functional (Figure S2A), but allowed quorum sensing across most of the full dynamic range of auxin concentration sensing (Figures 3B; Figure S2B). Cells responded similarly to cell density across different culture media volumes, indicating quorum sensing responded to cell density rather than absolute cell number (Figure 3C). Together, these results establish that Sender-Receiver cells can sense their own population density in two ways: using NAM without PIN2, or using either precursor with PIN2.

**Figure 3:**
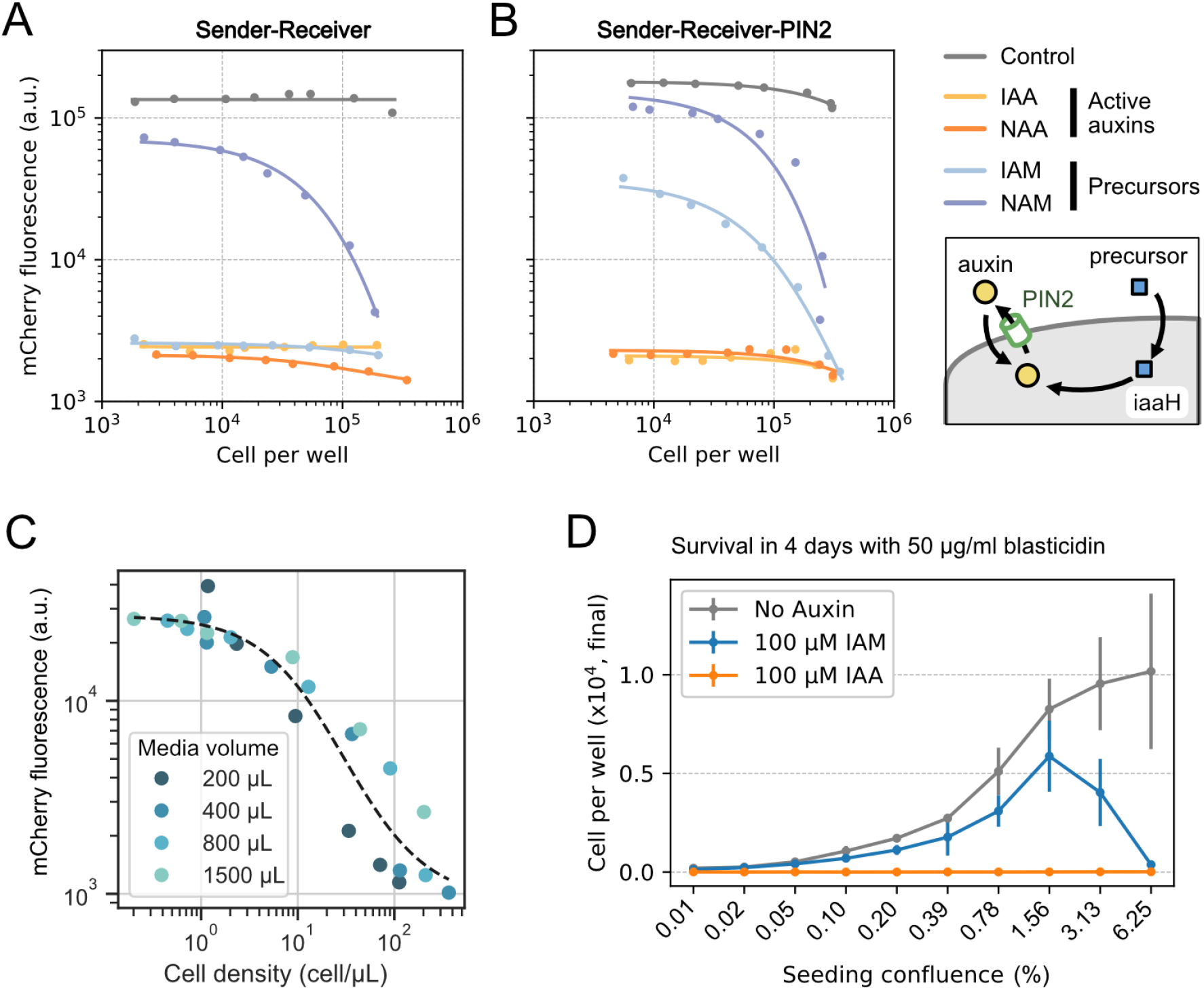
Sender-Receiver cells sense their population density and regulate survival accordingly. (A-B) Sender-Receiver and Sender-Receiver-PIN2 cells perform quorum sensing. For both plots, cells were seeded at different densities and induction conditions, with either of the two auxins (saturating signaling) or their precursors (to allow quorum sensing). mCherry fluorescence was assayed after two days as a reporter of auxin sensing. Inset: Cells in B express the transporter PIN2, which actively exports auxin. Data were fitted onto an inverted Michaelis-Menten’s function on log scale (see Figure S2B for fitted parameters, Method). (C) Sender-Receiver-PIN2 cells sense population size per unit volume. Cells were grown for two days at 6 different densities for each media volume, and cultured on a rocker for better mixing. (D) Sender-Receiver-PIN2 triggers cell death at high population density. Cells were seeded at different confluence levels, grown for four days in media with 50 μg/ml blasticidin and IAM, IAA, or no auxin. Error bars indicate standard deviation from triplicates.

### The Paradaux cell line implements paradoxical population control

Linking quorum sensing to cell survival could enable cell population control (Figure 1A, right). To test this possibility, we seeded cells at different densities in media containing blasticidin to make cell growth dependent on BlastR levels, as well as the precursor IAM. After 4 days, cells seeded at high, but not low, densities exhibited reduced cell numbers compared to cells plated at the same density without IAM or IAA (Figure 3D). These results demonstrate density-dependent control of cell survival.

This negative autoregulatory feedback loop resembles a similar design in bacterial systems, where a quorum sensing signal induces cell killing. The bacterial circuit was found to be susceptible to “cheater” mutations that allowed cells to escape control and grow to the limit of environmental capacity (Balagaddé *et al*., 2005). For adherent mammalian cells, environmental capacity is limited by the surface area of the culture plate. We therefore define an escape event as cells growing to confluency (covering the full surface of the plate) and remaining confluent for the duration of the experiment. When we cultured the Sender-Receiver-PIN2 cell line in media containing blasticidin and IAM to activate the circuit, we observed escape after 16 days of culture (Figure S2C, Movie S1), which reflects the acquisition of a “cheating” phenotype, with which cells are able to proliferate in high auxin (IAA) and blasticidin (Figure S2D). These results show that simple negative feedback population control circuits in mammalian cells, like their bacterial counterparts, are susceptible to selection pressure for cheater mutations.

A “paradoxical” circuit architecture, in which a quorum sensing signal either stimulates or inhibits both cell proliferation and cell death, can make population control more mutationally robust by counterselecting against cheaters that lose signal sensing (Karin and Alon, 2017). In the paradoxical circuits, positive net growth can occur only at signal concentrations lying between two non-zero bounds, similar to the Allee effect in ecology (Courchamp, Berec and Gascoigne, 2008) (Figure 4A, blue regions). By comparison, the simpler, non-paradoxical negative feedback architecture, in which signals only down-regulate growth, exhibits positive net growth at all signal concentrations below a single upper bound (Figure 4A, lower left). While both circuits exhibit stable fixed points at the maximum of their positive growth zones, they respond differently to cheater mutations that reduce auxin sensitivity (the predominant cheater phenotype observed in Figure S2C and S2D). Specifically, with negative feedback alone, such cheater mutations extend the regime of positive net growth to higher signal concentrations and cell densities, providing a growth advantage over non-mutant cells (Figure 4A, lower right). By contrast, in the paradoxical circuit, the same mutation would cause signal sensing to drop below the lower bound, activating the cell death arm of the circuit (Figure 4A upper right). In this way, the paradoxical circuit design should suppress escapes by cheater mutations that reduce or eliminate signal sensing.

**Figure 4:**
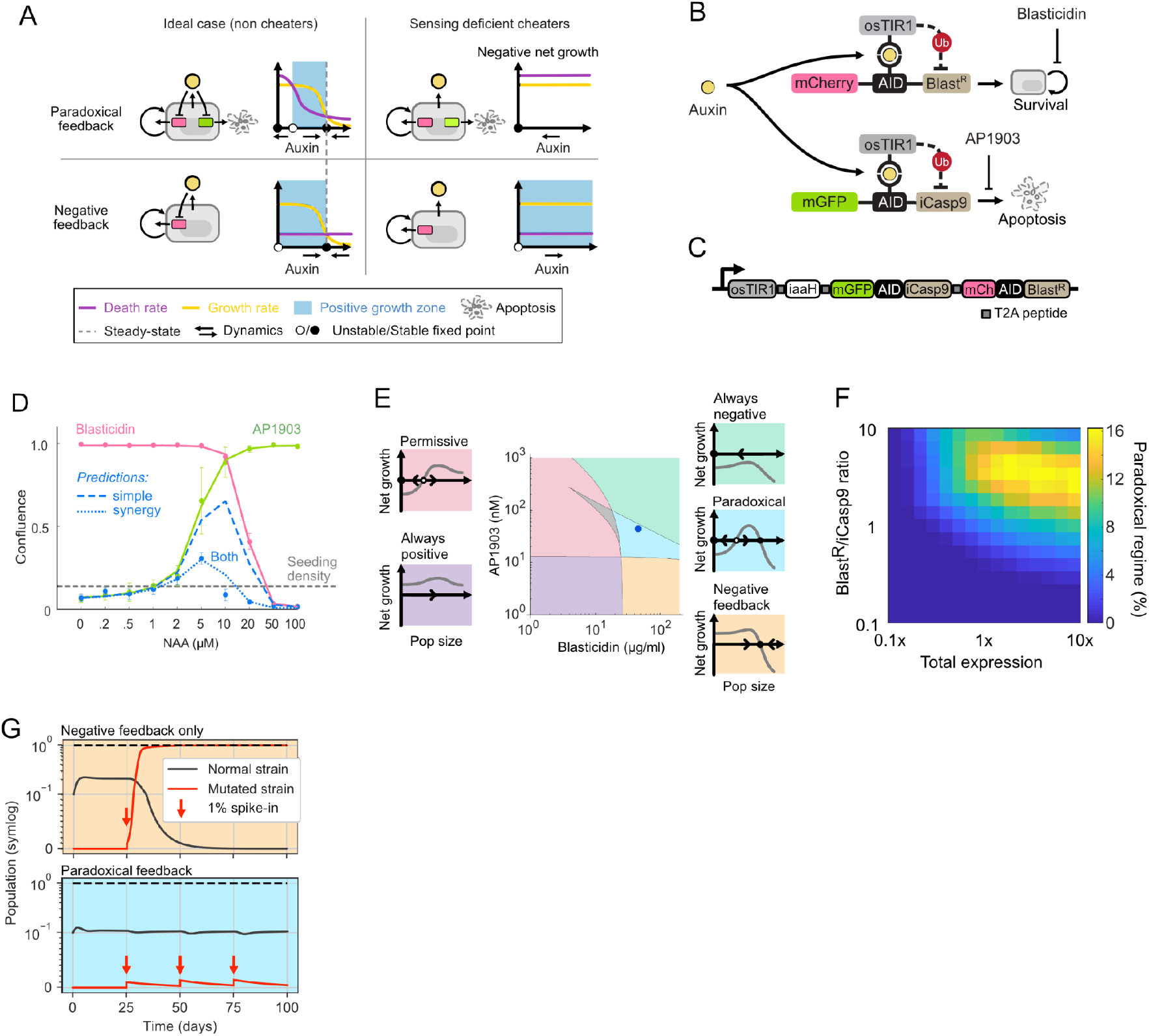
Paradoxical architecture reduces susceptibility to cheater mutations. (A) In the paradoxical architecture, the same signal inhibits growth (red pathway) and death (green pathway). This can produce a window of auxin concentrations leading to positive net growth (light blue region). Without mutation, the paradoxical and negative feedback circuits operate similarly around a stable equilibrium point of large population size (solid black dots, left panels). Mutations that eliminate sensing make both death and growth independent of auxin concentration (right panels), which selects against mutations in the paradoxical circuit due to negative net growth. (B) In the paradoxical circuit implementation, auxin regulates growth through BlastR-(upper path), and also regulates apoptosis via iCasp9 (lower path), each with distinct fluorescent protein readouts and a small molecule (blasticidin and AP1903) as a control switch. (C) The full paradoxical circuit can be encoded as a single open reading frame, with distinct proteins separated by T2A peptides (grey squares). (D) Paradaux cells respond to auxin in a biphasic manner. Cells were seeded at about 1/8 confluence (grey dashed line) and pretreated with NAA for one day, then treated with combinations of NAA, blasticidin (20 μg/ml) and AP1903 (50 nM) for another 3 days, and imaged. Pink, green, and blue dots show the mean and standard deviation of three replicates in the presence of AP1903, blasticidin, or both, respectively. Solid red and green lines indicate fits of these data to the model. Purple dashed lines indicate predictions for the fully operational circuit based on the green and red curves. Dotted purple line is the model prediction when the synergy term is included. (E) Different classes of behavior can occur in different parameter regimes. We simulated auxin-dependent growth in different parameter regimes and identified five distinct regimes (indicated schematically as insets). Using this classification, we numerically analyzed and sorted growth curves for each concentration of blasticidin and AP1903 (central plot). The blue dot indicates the blasticidin and AP1903 concentrations used in the time-lapse movie analysis (Figure 5). The grey region indicates curves that could not be classified into one of these categories (0.68% of total, see Figure S4F and STAR Methods). (F). For each expression level, we analyzed the percent of blasticidin-AP1903 concentrations that generate paradoxical behavior, similar to panel E. Optimizing the expression and ratio of BlastR and iCasp9 can widen the paradoxical regime. (G) Dynamic simulations show the Paradoxical Control circuit provides evolutionary robustness. For the negative feedback systemic and *γ_C_* are *γ_syn_* set to zero. Mutated strains were simulated with auxin (*A*) fixed to zero, representing sensing deficient mutations. On day 25, 50, 75 and 100, mutant cells (1% of the population cap) were introduced into the system. These mutants take over in the negative feedback circuit (top) but not the paradoxical circuit (bottom). Dashed line indicates carrying capacity.

We designed a paradoxical circuit in which auxin represses both proliferation and death, a configuration that is well-suited to the inhibitory nature of auxin-dependent protein degradation (Figure 4A, upper left). In addition to auxin regulation of BlastR, we added a parallel regulatory pathway in which auxin negatively regulates cell death by inducing degradation of iCasp9, an ectopically expressed master regulator of apoptosis (caspase 9) activated by the small molecular dimerizer AP1903 (Figure 4B) (Straathof *et al*., 2005; Di Stasi *et al*., 2011). We added an AID domain in iCasp9 to provide auxin regulation, and fused it to the monomeric GFP variant mGFPmut3, to allow direct readout of its concentration (Landgraf *et al*., 2012) (Figure 4B). This design allows one to operate the same cell line in three regimes, depending on what combinations of blasticidin and AP1903 are added to the medium. With only blasticidin, the circuit operates in the pure negative feedback regime; with blasticidin and AP1903, it operates in the paradoxical regime; and with neither inducer it has unregulated growth. Because it implements paradoxical regulation through auxin, we dubbed this circuit “Paradaux.”

To encode the Paradaux circuit, we designed a single multi-protein construct expressing osTIR1, the auxin-regulated iCasp9 system, and the auxin-regulated BlastR construct described above (Figure 4C). To eliminate one potential mechanism for evolutionary escape, we positioned BlastR at the C-terminal end of the construct, so that premature stop codon mutations would deactivate BlastR, decreasing survival. We integrated the construct to create stable monoclonal CHO-K1 cell lines for further analysis (Method). Because this line lacks PIN2, we used NAM/NAA rather than IAM/IAA for all Paradaux experiments. Survival of these cells increased monotonically with auxin in the presence of AP1903 and decreased with auxin in the presence of blasticidin, demonstrating that both branches of the Paradaux circuit were individually functional (Figure 4D, red and green lines). (An additional monoclonal line with an independent integration of the same circuit is shown in Figure S3A). Further, including both blasticidin and AP1903 produced a biphasic survival curve (Figure 4D, blue dots), the key requirement for paradoxical population control.

### Mathematical modeling identifies parameter regimes required for paradoxical control

To identify parameter regimes, including concentrations of blasticidin and AP1903, that optimize its population control capabilities, we developed a mathematical model of the Paradaux circuit. Assuming rapid intracellular-extracellular auxin equilibration, as observed for NAA (Figure 3A), and timescale separation between fast intracellular dynamics and the slower cell population dynamics (Figure S3B, STAR Methods), we derived an approximate model based on two differential equations. The first represents the extracellular auxin concentration shared by all cells, denoted *A* The second describes the cell population size, denoted *N*, which ranges from 0 to the environmental carrying capacity, normalized as 1 (STAR Methods).

We assume auxin is produced at a constant rate per cell, λ_*A*_, and diluted by periodic media changes, approximated as a continuous process with rate constant *δ*_*A*_:

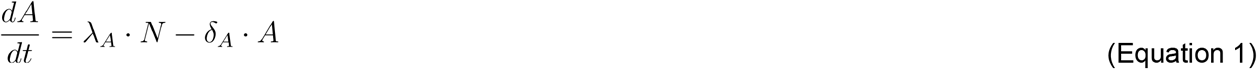

We also assume cell growth can be described by a generalized logistic function (Richards, 1959) (Figure S3C and STAR Methods), modified to incorporate the effects of blasticidin, *B*, and iCasp9, *I*, on growth:

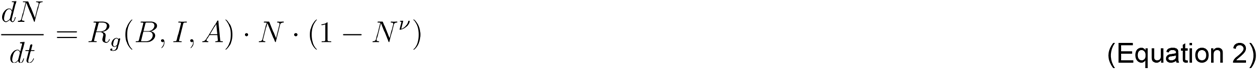

Here, *R_g_* denotes the cellular growth rate, a function that can be written as a sum of two Hill-like terms representing the combined auxin dependent effects of blasticidin and iCasp9 on cell survival (See Equations S13, S14, and S15 for the exact form). The exponent *v* is the non-linear correction parameter in the generalized logistic growth function (Richards, 1959).

To constrain the effective biochemical parameter values in the model, we incorporated experimentally measured values of the sensitivity of AID-tagged proteins to auxin, the unperturbed cell growth rate, the auxin secretion rate, and the non-linear growth correction parameter (Figure 1C; Figures S3C and S3D). Remaining parameters were fit using the auxin-dependent survival rates measured with either AP1903 or blasticidin (Figure 4D, red and green line). The model initially overestimated actual growth rates when both arms of the circuit were simultaneously active (Figures 4D, dashed purple lines), possibly due to previously reported synergy between apoptosis and blasticidin-dependent translational inhibition (Holcik and Sonenberg, 2005). We therefore added a phenomenological synergistic interaction term to the growth rate expression (Equation S17 and STAR Methods; Table S1, parameter set 1; Figure 4D, dotted line). Finally, we checked that experimental parameter values expected to be independent of integration site, such as maximum cell death and growth rates, as well as synergy and Hill coefficients, agreed, within ~2-fold, with those measured for a second Paradaux monoclonal cell line with an independent integration of the circuit construct (Figure S3A; Table S1, parameter set 2).

Before initiating challenging long-term analysis of population control, we used the model to systematically scan for AP1903 and blasticidin concentrations likely to favor paradoxical population control. For each pair of AP1903 and blasticidin concentrations, we classified the dependence of cell survival on auxin into one of five qualitatively distinct behaviors: (1) positive net cell growth across all auxin concentrations (Figure 4E, purple background, “uncontrolled growth” in Figure 5); (2) negative net cell growth across all auxin concentrations (Figure 4E, green background); (3) permissive growth, in which cells proliferate only beyond a minimum auxin concentration (Figure 4E, pink background); (4) simple negative feedback control, analogous to the Sender-Receiver behavior (Figure 4E, yellow background, “negative feedback in Figure 5”); and (5) paradoxical control (Figure 4E, blue background, “paradoxical feedback in Figure 5”). Only the negative and paradoxical feedback regimes produce a non-zero, stable fixed point, allowing population control. The desired paradoxical regime occurred in a window of blasticidin and AP1903 concentrations centered around 50 μg/ml and 50 nM, respectively (Figure 4E). Further, this paradoxical window could be enlarged by optimizing the expression levels of iCasp9 and BlastR (Figure 4F). Within the paradoxical regime, the circuit produced the expected bistability of population size (Figure S3E) and robustness to mutations that eliminate auxin sensing, simulated by setting sensed auxin to zero (Figure 4G).

**Figure 5:**
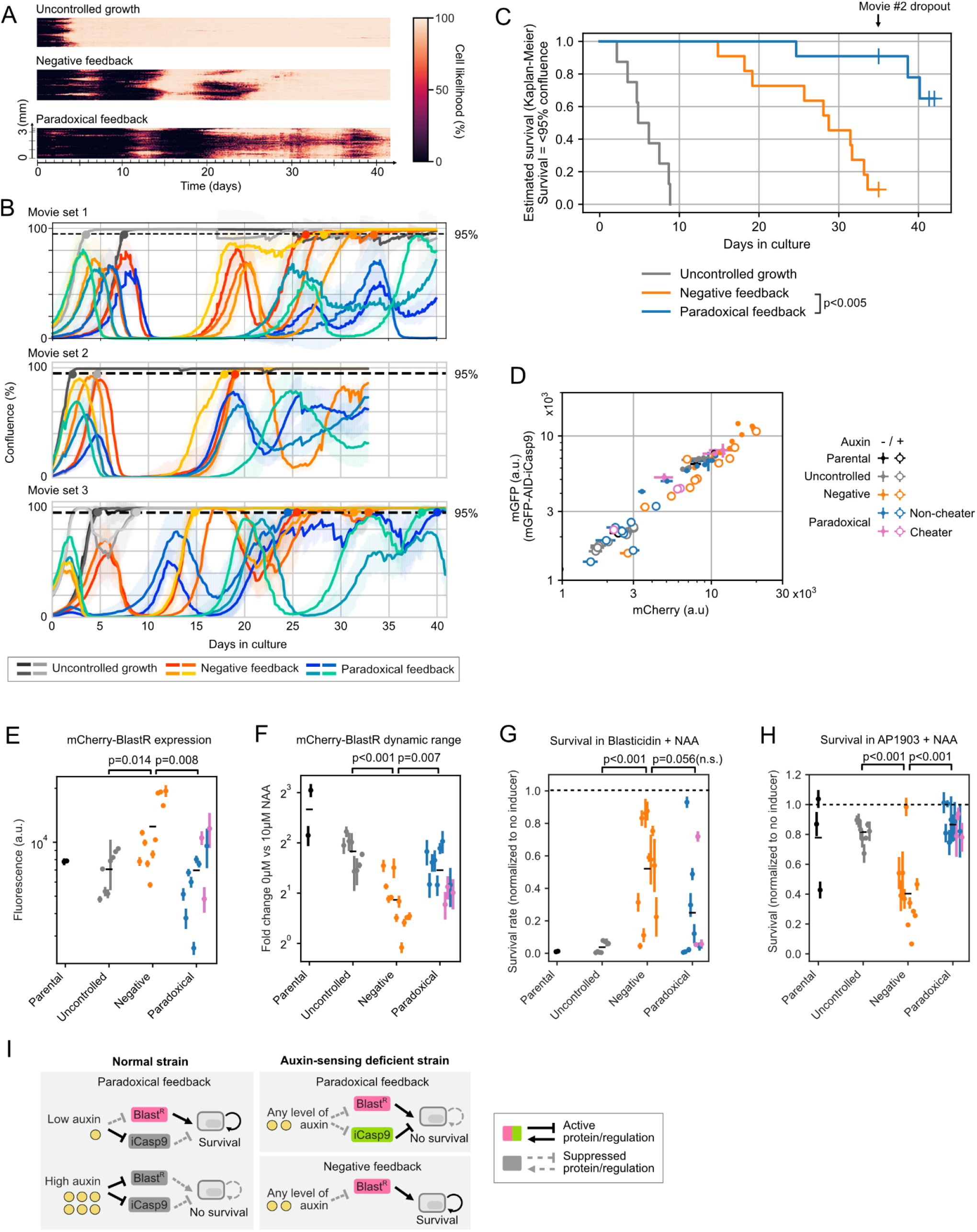
The paradoxical circuit allows mutationally robust population control. (A) Composite kymograph of long-term cultures with no control (100 μM NAM; upper panel), negative feedback (100 μM NAM and 50 μg/ml blasticidin; middle panel), or paradoxical feedback (100 μM NAM, 50 μg/ml blasticidin, and 50 nM AP1903; lower panel), from movie set 3. For visualization, the images were analyzed using ilastik and 30-pixel-wide strips from each timepoint were combined to make the kymograph. (B) Population dynamics for the three movie sets conditions reveal delayed mutational escape for the paradoxical circuit (shaded envelopes represent the standard deviation across 12, 25, and 36 stage positions, for each movie set respectively). Solid dots indicate escape events, defined by cells exceeding 95% confluency and not returning below that threshold for the duration of the movie. Black arrows indicate late cheating isolates that are similarly denoted with arrows in (G). (C) Kaplan-Meier estimate (Kishore, Goel and Khanna, 2010) of survival (no mutant escape) for movies in (B). Samples in movie set 2 that ended earlier were treated as dropouts. The samples under paradoxical feedback retain population control significantly longer than those under negative feedback (p<0.005, log-rank test). (D-F) Isolates were treated with 10 μM NAA, or nothing, for two days prior to flow cytometry assay for mCherry and mGFP fluorescence, co-expressed with BlastR and iCasp9 respectively. Bars represent standard deviation from triplicates (D and E), or bootstrapping (F, n=9). (D) Isolates from all long term cultures show correlated BlastR and iCasp9 expressions. (E) Isolates from negative feedback, but not paradoxical feedback, showed upregulation in BlastR, compared to control. (F) Isolates from negative feedback showed diminished dynamic range compared to both control and paradoxical groups. (G and H) BlastR and iCasp9 expression levels affect survival rates across different conditions. Cells were seeded in 96-well imaging plates with IAA or standard media for one day, and the second drug, blasticidin (G) or AP1903 (H), was added. Samples were then imaged to estimate confluency at day 4. For each isolate, survival rates were normalized to the group with no second drug. The black arrows in (G) highlight the isolates that cheated at the final days in movie set 3, black arrows in (B). Values and errors were calculated by bootstrapping (n=6, Methods). (I) How the paradoxical architecture, but not negative feedback, eliminates cells that lose the ability to sense auxin (schematic).

Additionally, we explored the effect of the 2-3 day time delay required for blasticidin to kill sensitive cells (Sato *et al*., 2012) (Figure S3F). Time delays in negative feedback loops are known to produce oscillations under some conditions (Elowitz and Leibler, 2000; Potvin-Trottier *et al*., 2016). In simulations of the paradoxical circuit, inclusion of a time delay led to oscillations of population density with periods of 2 weeks or more depending on the value of the delay parameter, *τ* (Equation S20). A bifurcation (Martinez-Corral *et al*., 2018) between damped and sustained oscillations occurred at approximately *τ* = 48 *hrs* (Movie S2). Taken together, these results provided insight into parameter dependence and expected dynamics of the circuit, and identified specific AP1903 and blasticidin concentrations for long timescale analysis of the experimental circuit.

### Paradoxical circuits extend the duration of population control

To experimentally analyze population control, we continuously monitored cultures of the Paradaux cell line using time-lapse movies (Figure 5B). We performed three sets of time-lapse experiments, the longest of which lasted for 42 days (an arbitrary time scale constrained by technical limitations). Cells were cultured in regimes of uncontrolled growth (no blasticin or AP1903, the “always positive” regime in Figure 4E), negative feedback (blasticidin added), or paradoxical feedback (both blasticidin and AP1903). In the uncontrolled regime, cells grew to and remained at full confluence for the duration of the movie (Figures 5A, Supplementary movie S3). In the latter two regimes, population dynamics exhibited oscillations, consistent with models incorporating time delays for blasticidin-dependent cell killing (Figure S3F).

The negative feedback regime limited population size for 1 or 2 oscillatory periods (~20-30 days), after which cells escaped control (exceeded >95% confluence) for the remainder of the movie (at least 10 subsequent days, Figure 5B). This behavior is consistent with the experimentally observed susceptibility of simple negative feedback control to escape mutants (Figure S2C and S2D), and simulated responses to introduction of sensing mutants (Figure 4G). By contrast, in the paradoxical regime, cultures exhibited oscillations in population density, but remained at sub-saturating densities for significantly longer (p<0.005, Kaplan-Meier estimate, log-rank test (Kishore, Goel and Khanna, 2010)). In fact, over half of the cultures remained sub-confluent for the full duration of the movies (Figure 5A-C; Supplementary movie S3). More specifically, across the three movie sets and the 11 individual cultures in the paradoxical group, only one culture escaped control before the 32 day mark (Figure 5A, Movie set 3). Together, these results demonstrate that the paradoxical control circuit reproducibly extends the duration of population control.

To gain insight into how cells evolved during long term culture, we isolated cells at the end of each of the 30 individual movies, passaged them in standard media, and assayed their behaviors under different conditions (Figure 5D-H). The fluorescence of mCherry and mGFP, which are co-expressed with BlastR and iCasp9, respectively, varied substantially across all isolates, reflecting independently acquired changes in the expression of circuit components, rather than amplification of a common founder mutation already present in the parental cell line. However, variation of the two reporters was strongly correlated expression across isolates, suggesting that mutations or other adaptations occurred upstream of both circuit arms (Figure 5D).

The two population control conditions produced different effects on circuit component expression and phenotypic behavior. In cells selected under negative feedback conditions, basal BlastR expression was upregulated, on average, by 73%, and showed 49% less responsiveness to auxin compared to the uncontrolled group (Figure 5E and 5F; p = 0.014 and p<0.001, respectively). By contrast, isolates from the paradoxical feedback conditions showed significantly lower basal BlastR expression (p = 0.008) and retained a larger dynamic range of BlastR regulation compared to negative feedback isolates (p = 0.007). These differences in BlastR regulation were also reflected in cell survival (Figure 5F and 5G). When cultured in a combination of blasticidin and NAA, mimicking high cell density, negative feedback isolates exhibited increased survival compared to the uncontrolled group (p<0.001 Figures 5G). While individual paradoxical isolates also exhibited elevated survival, the increase compared to the uncontrolled condition was not significant at the group level (p=0.085). We also note that the two isolates which were classified as cheaters in the final days of movie set 3 (Figure 5B, black arrows) were unable to survive in this condition (Figure 5G, pink dots with arrows), suggesting they could potentially have returned to lower cell density had the movie duration been longer. Together, these results suggest that paradoxical conditions preserved more of the original BlastR regulation and function compared to negative feedback conditions.

The paradoxical architecture is designed to suppress sensing deficient mutations that reduce the responsiveness of both arms of the circuit to auxin. More specifically, cells that lose auxin sensing should no longer degrade BlastR (Figure 5I, lower right), and therefore obtain a growth advantage. By contrast, in the paradoxical circuit, such sensing deficient cells would also be unable to degrade iCasp9, leading to their elimination (Figure 5I, upper right). Thus, sensing deficient isolates from the negative feedback condition should become susceptible to killing through the iCasp9 arm, even in the presence of auxin. In fact, isolates from negative feedback conditions were sensitive to the combination of AP1903 and auxin, indicating that they had acquired the potential to be counter-selected under paradoxical conditions (Figure 5H, p<0.001). Thus, cells with reduced sensing gain a growth advantage in negative feedback conditions but are counter-selected under paradoxical conditions, consistent with the principle of paradoxical control.

To better understand the heritable changes that drive adaptation during long term culture, we sequenced the circuit from endpoint isolates from the first two movie sets. Several mutations appeared at elevated frequencies, occurring either in the AID domain fused to BlastR, consistent with positive selection through reduction of AID activity, or in osTIR1, near the 5’ end of the circuit transcript, potentially reducing expression of all circuit components (Figure S4A; Table S2). However, in most isolates, no mutations occurred at significantly elevated frequencies, suggesting that most relevant adaptations (observed in Figure 5D–5H) lie outside the circuit itself.

Therefore, we performed RNA-Seq expression analysis to investigate global transcriptional changes. Expression profiles from isolates cultured in the same conditions generally clustered together, suggesting that each operating mode of the circuit selects for a distinct set of gene expression changes (Figure S4B). In addition, solates from the uncontrolled growth conditions clustered with the parental cells, as expected. Pathway enrichment analysis further revealed that ribosome components, the primary targets of blasticidin, were significantly upregulated in both negative feedback and paradoxical conditions (Figure S4C). Proteasome components were up-regulated only in negative feedback conditions (Figures S4C and 4D), consistent with opposite selection pressures on proteasome activity generated by auxin regulation of BlastR and iCasp9 in the paradoxical condition.

Taken together, these results indicate that diminished sensitivity to auxin provides the dominant growth advantage in the negative feedback condition (Figure 5E and 5F), but is counter-selected in the paradoxical condition, where it triggers activation of iCasp9 (Figure 5H). By contrast, in the paradoxical condition, escape paths are more limited, accounting for the extended duration of population control.

## Discussion

Natural cytokine-based control circuits allow cells to regulate their own population dynamics, as well as those of other cell types. Synthetic circuits could provide analogous capabilities. To this end, we engineered simultaneous production and sensing of the plant hormone auxin in mammalian cells, and coupled it to genes controlling cell proliferation and death. The enzymes iaaH and iaaM, together with PIN2, allow cells to produce and export auxin. osTIR1 together with AID domains provide a simple, direct means of sensing auxin and coupling it to arbitrary protein targets (Figures 1–2). These components thus provide a long-range private communication channel, and enable the foundational property of quorum sensing (Figure 3). Coupling quorum sensing to cell survival opens up the possibility of creating population control circuits, provoking the question of what circuit architectures can provide robust, long-term control. Consistent with previous work (Karin and Alon, 2017), mathematical modeling showed that a paradoxical architecture, in which auxin inhibits survival mediated by BlastR and killing by iCasp9, can generate a range of qualitatively different behaviors and, in some regimes, suppress cheaters (Figure 4E and 4G). To experimentally realize this capability, we constructed the “Paradaux” circuit, and compared its operation in three distinct regimes—uncontrolled growth, negative feedback, and paradoxical—by using media with different combinations of blasticidin and AP1903 (Figure 4B). Long-term culturing for up to 43 days revealed that while both negative and paradoxical feedback architectures can limit cell population size for weeks, the former is more susceptible to escape by sensing-deficient cheaters (Figures 5, S2B, and S4). By contrast, the paradoxical design suppressed these cheaters, as predicted theoretically (Figures 4 and 5) (Karin and Alon, 2017), and provided more robust population control for the conditions and timescales explored here.

In addition to quorum sensing and population control, the auxin cell-cell communication system opens up the possibility of engineering more complex multi-cell type communication and control systems. For example, to record the relative distance between two groups of cells, one could engineer receivers that permanently activate in response to auxin secreted by a second cell type. Alternatively, one could take advantage of the two-step nature of auxin biosynthesis (Figures S1A and S1C) by separating the steps into distinct cells, thereby enabling a proximity-dependent AND gate. More complex cytokine-like, multi-channel circuits could be achieved by combining the auxin with additional diffusible, orthogonal signals. This approach could enable synthetic bidirectional signaling (Zhou *et al*., 2018) or even Turing-like spatial pattern formation (Turing, 1952). Candidate molecules for additional channels include the plant hormones abscisic acid and gibberellin, both of which have partially mapped biosynthetic pathways (Early and Martin, 1990; Mashiguchi *et al*., 2011; Finkelstein, 2013) and were previously engineered to induce protein-protein interactions, enabling them to regulate transcription or protein localization in mammalian cells (Liang, Ho and Crabtree, 2011; Miyamoto *et al*., 2012). The plant IP-CRE1 cytokinin system, which has been successfully ported to yeast, uses a plant-specific hormone as a diffusible signal to induce phosphorylation and further transcriptional activation, providing an additional potential signaling system (Chen and Weiss, 2005). Bacterial autoinducers and synthetic proteins also provide attractive candidates for additional orthogonal signaling channels (Hong *et al*., 2012; Daringer *et al*., 2014; Morsut *et al*., 2016).

With these developments, private communication channels, population sensing, and population control could improve engineered cell therapies by allowing cells to coordinate their responses and localize activities at target sites. Previously, several studies attempted to activate latent natural signaling abilities to trigger coordinated multicellular responses. For example, to locally stimulate immune function, T cells were engineered to secrete IL-12 and IL-18 upon tumor infiltration (Chmielewski *et al.*, 2011; Hu *et al*., 2017). Similarly, secretion of bispecific T-cell engagers (BiTEs) by engineered T cells was shown to guide bystander T-cells to attack at the cancer site, successfully improving infiltration and reducing toxicity caused by normal tissue expressing the target (Choi *et al*., 2019). Private auxin-based communication channels would complement these approaches by allowing engineered cells to not only specifically sense and limit their own local population size but also to enable conditional activation only beyond a minimum density. We therefore anticipate the incorporation of synthetic population control systems in future generations of engineered cell therapies.

Further work could improve the circuits presented here and circumvent their limitations.The Paradaux cell line requires addition of the auxin precursors IAM or NAM for auxin synthesis. Though this provides a good external control method for our experimental setup, it might be suboptimal in therapeutic scenarios. For these applications, complete biosynthesis of IAA can be achieved by expressing tryptophan 2-monooxygenase (iaaM), in conjunction with iaaH (Figure S1A and S1C). The use of blasticidin and its auxin-regulated resistance gene for growth control is not ideal, since antibiotics have complex effects on the cell, must be added to the media, and broadly affect all cells exposed to the drug, making them inappropriate for cell therapy applications. Enzyme-prodrug systems could provide an alternative approach with fewer non-specific effects on host cells (Sharrock *et al*., no date; Trask et al., 2000). Alternatively, a more generalizable, cell-autonomous system could be achieved by coupling auxin to a cell-cycle regulator, such as Cdk1, 2 or 3 (Satyanarayana and Kaldis, 2009), or other genes (Harborth *et al*., 2001) essential for survival and/or cell cycle progression. Due to the inherent time delays within the feedback loop, the Paradaux circuit exhibits oscillatory behaviors (Figure S3F), similar to earlier synthetic population control circuits in bacteria (Balagadde, 2005). Reducing feedback delays or implementing more sophisticated control systems (Chevalier *et al*., 2019) should facilitate non-oscillatory homeostatic dynamics. Finally, although the paradoxical control system successfully extended the duration of population control, cells nevertheless accumulated adaptive changes (Figure 5 and S4). Future work exploring even longer timescales should reveal how long the paradoxical design can extend the duration of control in the presence of strong selection pressure to subvert it.

## Acknowledgments

We thank Igor Antoshechkin, Vijaya Kumar and Jeff Park for technical assistance and advice; Xinying Ren and Leopold Green from Murray lab for discussion; Haley Larsen, Ke-Kuan Chow, Felix Horns, Duncan Chadly, Christina Su, Ronghui Zhu, Lucy Chong and other members of the Elowitz lab for critical feedback on the manuscript; and Uri Alon, Omer Karin, Siyu Chen and Matt Thomson for scientific input and advice. This work is based on work supported by the Defense Advanced Research Projects Agency under Contract No. HR0011-17-2-0008, by the National Institute of Health grant R01 MH116508, by the Allen Discovery Center program under Award No. UWSC10142, a Paul G. Allen Frontiers Group advised program of the Paul G. Allen Family Foundation, and by the Millard and Muriel Jacobs Genetics and Genomics Laboratory at California Institute of Technology. M.B.E is a Howard Hughes Medical Institute Investigator. A.L. Is supported by a fellowship from the Taiwanese Ministry of Education. The content of the information does not necessarily reflect the position or the policy of the Government, and no official endorsement should be inferred.

## Author contributions

Project direction, supervision, and funding: MWB, MBE, RMM; Design of research: YM, MWB, MBE; Experimental investigation: YM, MWB, JZ, AL; Data analysis: YM, MWB, MNM, MBE; Mathematical modeling: YM, MNM, MBE; Paper writing: YM, MWB, MWM, MBE.

## Supplemental figures and legends

**Figure S1:**
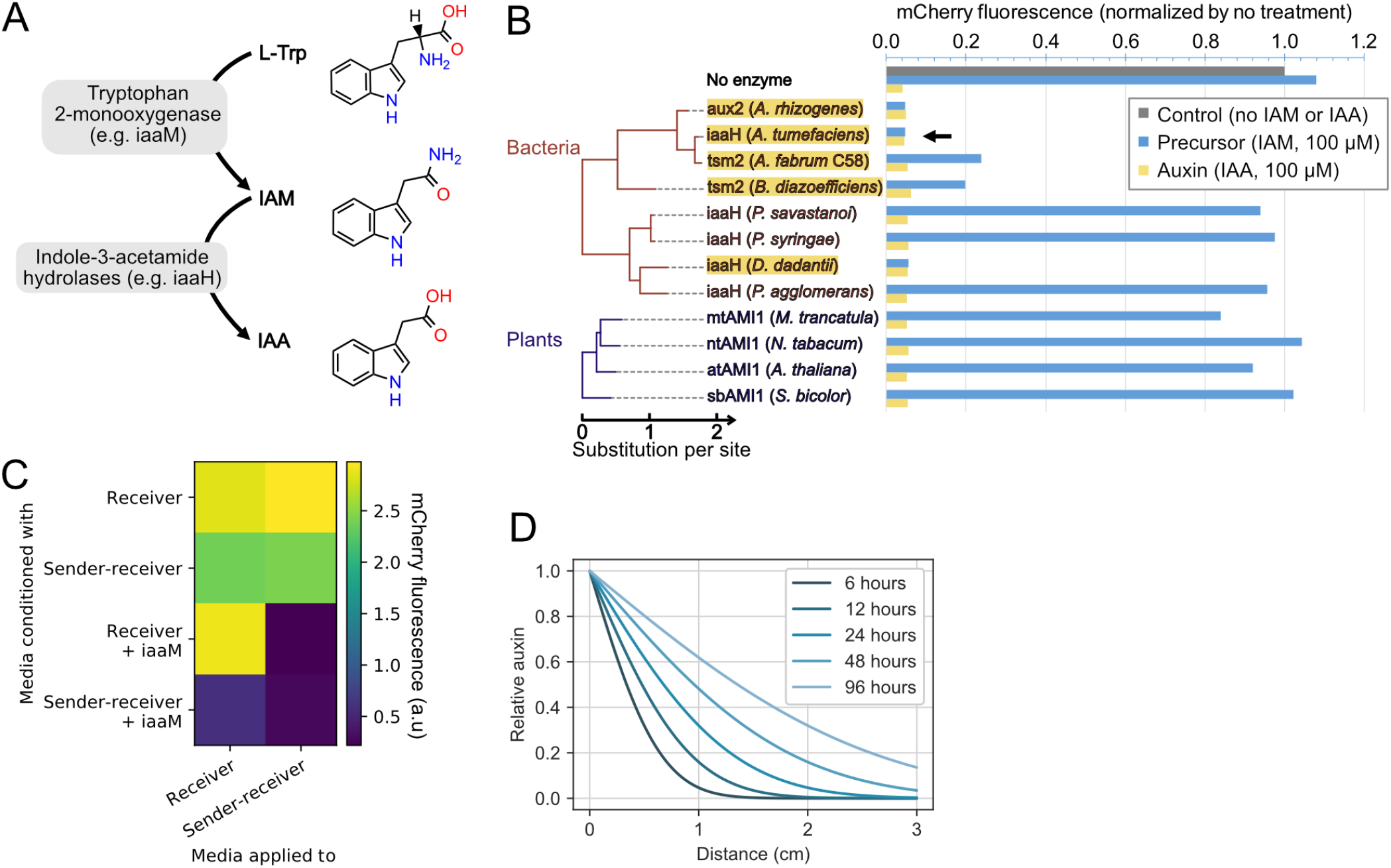
Expression of auxin-synthesizing enzymes enables cells to produce auxin, related to Figure 2. (A) IAA can be synthesized from L-tryptophan (L-Trp) in a two-step pathway. (B) Twelve candidate indole-3-acetamide hydrolases were screened for the ability to convert IAM to IAA. The phylogenetic tree was created with CLUSTAL-OMEGA (Sievers *et al*., 2011) and phyML3.0 (Guindon *et al*., 2010) using default settings. Candidate enzymes were transfected into Receiver cells, and auxin production was assessed by mCherry-AID degradation (Figure 2B) after 48 hours. The fluorescent results are normalized to the control without auxin. Candidates that could down-regulate the mCherry by over 50% are highlighted in yellow. iaaH from *A. tumefaciens* (black arrow) was selected for subsequent experiments. (C) Media conditioned by iaaM (from *P. savastanoi)* expressing cells contains substrate for auxin production by iaaH. Cells transfected with iaaM were used to condition the media for 48 hours, as shown in Figure 2C. Conditioned media was combined with an equal amount of fresh media and then applied to reporter cells for 48 hours before flow cytometry. (D) Auxin diffusion observed in Figure 2E matches theoretical predictions (STAR Methods).

**Figure S2:**
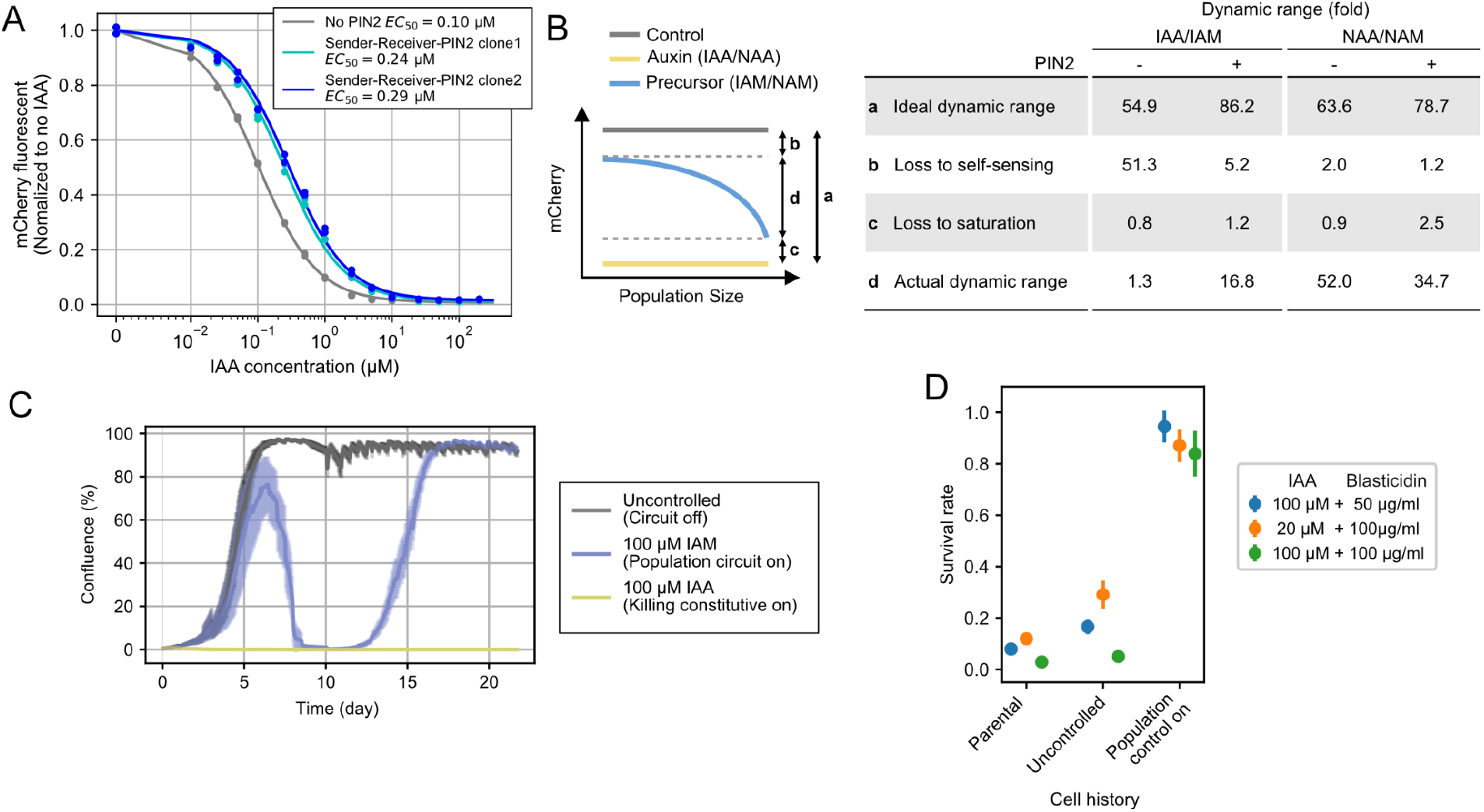
Sender-Receiver cells sense their population density andc regulate survival accordingly, related to Figure 3. (A) The right-shifted auxin response curve of PIN2 expressing cells is consistent with active export of IAA from the cells. Two PIN2-expressing cell lines were incubated with IAA for 48 hours and then assayed for mCherry fluorescence. The data were fitted with the same method in Figure 1C. (B) Quantitative analysis of Figure 3A and 3B shows that the expression of PIN2 rescues dynamic range (indicated by the double-sided arrows in the left panel) lost to self-sensing (STAR Methods). (C) A pilot experiment showed that the Sender-Receiver-PIN2 cells escape regulation after 15 days of continuous culture. Cells were seeded into a 24-well imaging plate with 50 μg/ml of blasticidin solely (Circuit off) or also with 100 μM IAA or IAM (Circuit on). Each trace reflects an average of 12 positions in a sample well. Shaded envelopes represent the standard deviation of the averages. (D) Cheater cells collected from the end of the movie in (C) have high survival rates when grown in IAA with blasticidin for 4 days as compared to parental cells. Cell counts measured with flow cytometry were normalized to matched cells grown without drugs. Bars represent standard deviation from triplicates.

**Figure S3:**
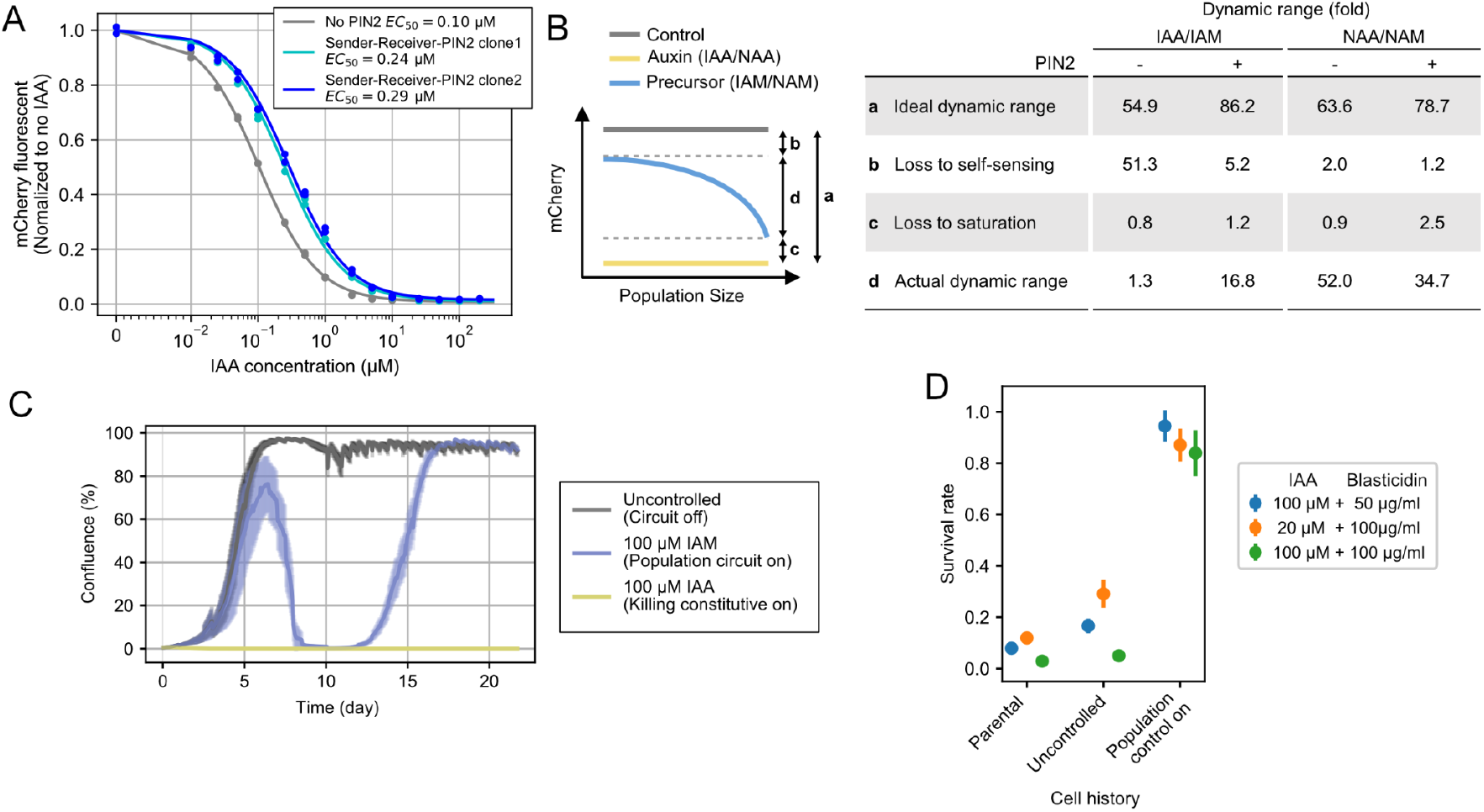
Model and parameter fitting of the paradoxical circuit, related to Figure 4 and Table S1. (A) A second Paradaux line responds to auxin in a biphasic manner. 4 nM of AP1903 and/or 50 μg/ml blasticidin were used; other parameters were the same as those used in Figure 4D. Curves are fitted directly from the survival data because the maximum growth rate is not available (Table S1). (B) The complete and reduced (approximate) versions of the model show similar dynamics, (overlap of black and cyan trajectories, see STAR Methods). (C) Generalized logistic growth (Equation 2) fits PC1’s growth curve better than a standard logistic growth curve. Data are from the first 120 hours of the uncontrolled growth sample in movie set 1, Figure 5B, seeded at 3000 cells per well. Data is smoothed with a Gaussian filter. Fitting results: *α* = 0.0395/*hr; v* = 2.26. (D) Calculation of auxin secretion rate (*λ_A_*). 2.09*μM*/106*cell · hr*. Sender-Receivers were seeded at 10,000 cells per well. After different durations (0 to 73.5 hours), conditioned media was collected to assay auxin concentration and cells were counted to estimate exponential growth rate. To determine the auxin concentration, the conditioned media was applied to Receivers at 1X, 0.5X, or 0.25X concentrations, and then Receiver fluorescence was compared to a standard curve. (E) Dynamic simulation of the system shows biphasic growth. Simulations were initiated at different cell densities. For each initial cell density, the initial auxin concentration was determined by setting Equation 1 to equal zero (equilibrium for auxin). (F) Simulation with delay (*τ*, Equation S20) incorporated show oscillated behaviours. Cells were seeded at 0.1, and the delay was applied to the blasticidin related response only. (G) Sub-sampling of the “unclassified” curves from Figure 4E shows that they are intermediate between “permissive” and “paradoxical” categories.

**Figure S4:**
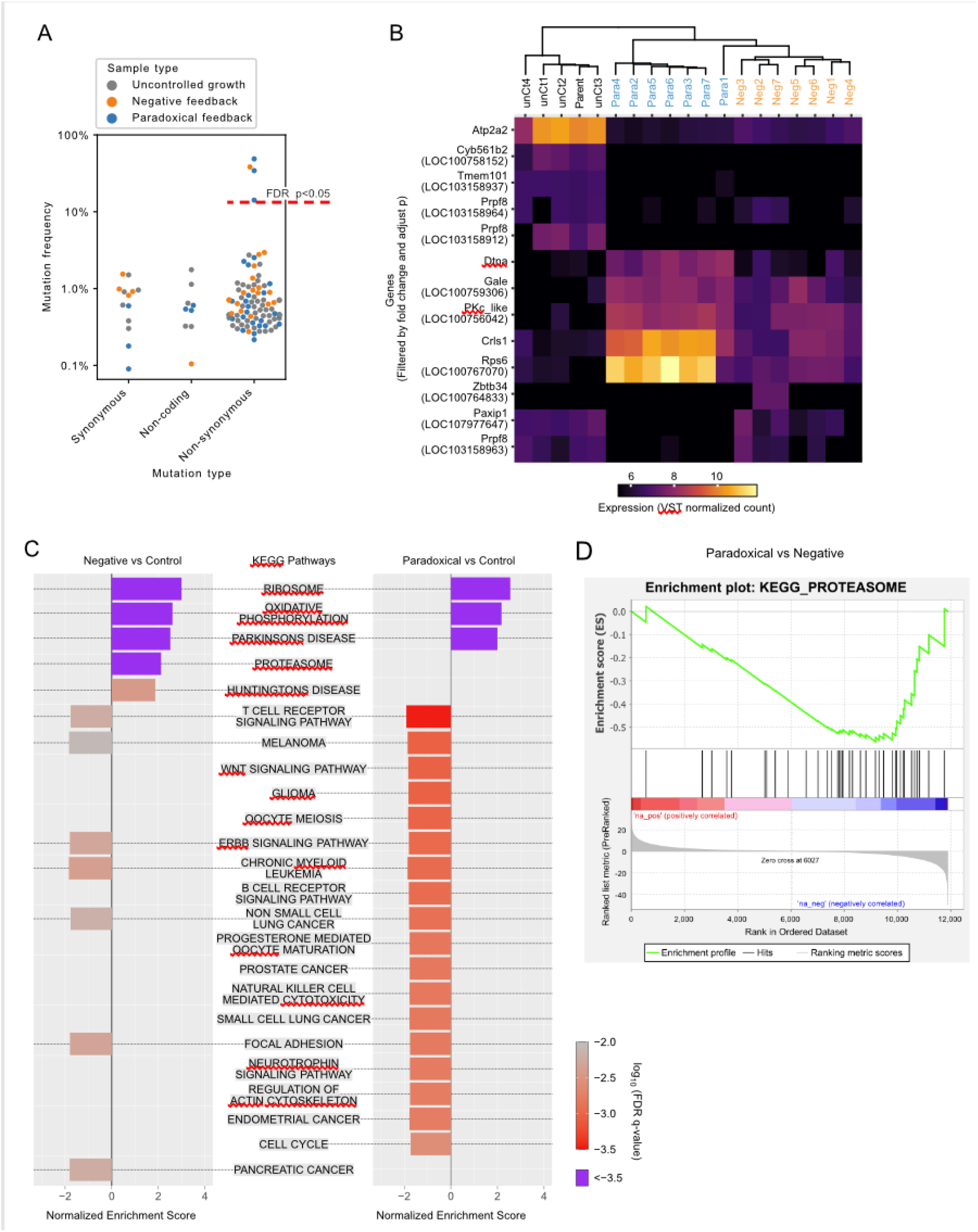
Targeted DNA and whole-genomic RNA sequencing reveal mutations and gene expression changes during long term culture, related to Figure 5 and Table S2. (A) Four mutations of significant positive selection were detected in the circuit component sequencing (details listed in Table S2). The integrated transgene was amplified and sequenced (Illumina). Mutations were detected by comparing reads of isolates to those of the parental line (chi-square test, Bonforroni corrected p<0.05, Method). The frequency threshold for positive selection (red dashed line) was determined as the 5th percentile (Bonferroni corrected) of the distribution of synonymous mutations, assumed to follow normal distribution. (B) Isolates that evolved in the same conditions clustered together, based on variance-stabilizing transformed (VST) read count across the whole transcriptome. Genes that are significantly differentially expressed between the groups (fold change > 32, adjusted p<0.01) are shown here (DESeq2, (Love, Huber and Anders, 2014)). Genes that lack annotation in RefSeq were manually annotated (Method), with the RefSeq number in parentheses. (C) Pathway enrichment analysis shows that the negative feedback and paradoxical feedback affect different sets of pathways. Enrichment analysis was performed with Gene Set Enrichment Analysis’ (GSEA) ranked method against MSigDB’s curated version of KEGG canonical pathways (Kanehisa *et al*., 2021). (D) The negative and paradoxical circuit design exert different evolutionary pressures on the proteasome pathway. Direct comparison of the KEGG proteasome pathway between the paradoxical and negative group was analyzed by GSEA, as shown with a standard GSEA plot. Here, blue (red) regions indicate genes whose expression is significantly greater (lower) in the negative feedback condition compared to the paradoxical condition.

## Supplemental Tables and legends

**Table S1:** List of parameters and fitted values, related to Figure 4 and S3. Fitted values are labeled as (1) or (2), in cases where the major Paradaux line (labeled as 1, Figure 4D), and the one used from confirming the model (labeled as 2, Figure S3C) were fit with different values. Dimensionless values are labeled as “DL” in the “Unit” column.

| Parameter | Description | Fitted value | Unit |
| --- | --- | --- | --- |
| $A$ | Production rate of auxin at 100% confluence | 4.18 | $\mu\text{M/hr}$ |
| $\delta_A$ | Dilution rate of auxin | 0.0289 | $\text{hr}^{-1}$ |
| $\nu$ | Generalized logistic growth coefficient | 2.69 | DL |
| $\alpha$ | Maximum natural cell proliferation rate | 0.0395(1); 0.0327(2) | $\text{hr}^{-1}$ |
| $\beta_B$ | Maximum net growth reduction by blasticidin | 0.0874(1); 0.147(2) | $\text{hr}^{-1}$ |
| $\beta_C$ | Maximum death rate induced by iCasp9 | 0.0451(1); 0.0375(2) | $\text{hr}^{-1}$ |
| $\beta_{syn}$ | The synergetic coefficient | 1.12(1); 1.70(2) | $\text{hr}^{-1}$ |
| $\kappa_A$ | Effective strength of auxin | 1.316 | $\mu\text{M}^{-1}$ |
| $\kappa_B$ | Effective strength of blasticidin | 0.045(1); 0.069(2) | $(\mu\text{g/ml})^{-1}$ |
| $P_R$ | Effective expression rate of BlastR | 1.77(1) ;0.716(2) | DL |
| $\kappa_I^{-n_2} \cdot P_C$ | Effective expression rate of iCasp9 | 8.54e-4(1);1.23(2) | DL |
| $n_1$ | Cooperativity of the blasticidin regulation | 2.26(1); 2.28(2) | DL |
| $n_2$ | Cooperativity of the iCasp9 regulation | 1.41(1); 1.67(2) | DL |

**Table S2:**
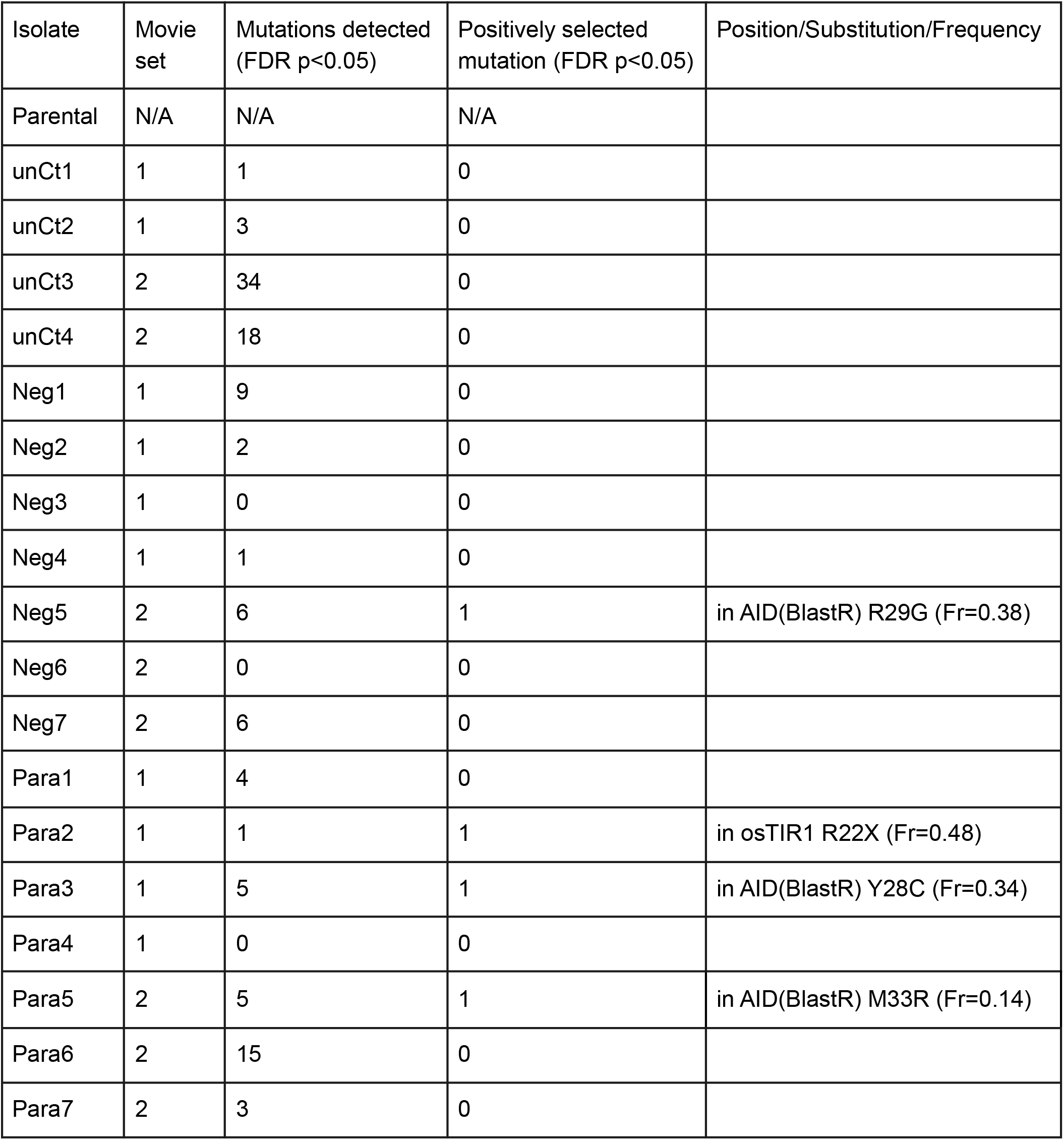
Details of the mutations detected in DNA sequencing results, related to Figure 5 and S4. Circuit components of isolates from movies set 1 and 2 were amplified and sequenced. For simplicity, uncontrolled growth, negative feedback and paradoxical feedback conditions are noted as unCt, Neg and Para, respectively, in the table. Four out of 18 sequenced isolates exhibited non-synonymous mutations at frequencies that were significantly elevated compared to those of synonymous mutations, consistent with positive selection (Bonferroni corrected, p<0.05, Figure S4A). Three of these (one in a negative feedback cheater isolate, the other two from non-cheating paradoxical isolates) squired mutations in the AID domain of the AID-BlastR fusion protein. Reduced activity of this AID domain would be expected to increase fitness under both paradoxical and negative feedback conditions. The fourth mutation, in a non-cheating paradoxical isolate, was a premature stop codon in osTIR1, near the 5’ end of the circuit transcript, which could potentially eliminate expression of all circuit components from the integrated copy in which it appears. (Because there are multiple genomic integrations of the circuit, any individual mutation could have a limited beneficial effect.)

## STAR Methods

### RESOURCE AVAILABILITY

#### Lead contact

Further information and requests for resources and reagents should be directed to and will be fulfilled by the Lead Contact, M. Elowitz.

#### Material availability

- Plasmids generated in this study have been deposited to Addgene, at https://www.addgene.org/plasmids/articles/28212030/
- Key cell lines will be available upon request.

#### Data and code availability

- The targeted DNA sequencing data have been deposited at NCBI Sequence Read Archive and are publicly available as of the date of publication. Accession numbers are listed in the key resources table.
- The bulk RNA sequencing data have been deposited at GEO and are publicly available as of the date of publication. Accession numbers are listed in the key resources table.
- The plasmid GenBank files, raw data, and processing/plotting scripts for generating the figures shown in this paper are available at data.caltech.edu. The DOI is listed in the key resources table.
- All original codes used for processing the images and the movies, and codes for mathematical modeling section (Figure 4 and S3) is available at github and is publicly available as of the date of publication. DOIs and links are listed in the key resources table.

### EXPERIMENTAL MODEL AND SUBJECT DETAILS

#### Tissue culture and cell lines

CHO-K1 (Hamster cells, RRID:CVCL_0214, ATCC Catalog No. CCL-61) cells and their derivatives were grown on tissue culture grade plastic plates (Thermo Scientific) in Alpha MEM Earle’s Salts (Irvine Scientifics), supplemented with 10% Tet System Approved FBS (ClonTech, or VWR), 100 U/ml penicillin, 100 μg/ml streptomycin, 0.292 mg/ml L-glutamine (GIBCO) or 1x GlutaMax (GIBCO). The complete media is filtered with 0.22 micron filters (Falcon).

For long-term culturing demonstrated in Figure 5, cells were seeded in 24-well TC-treated plates (total media 500 μL per well) with imaging-grade plastic bottoms (ibidi inc. #82406), and media was changed every 12 hours with one of the following methods: 1) adding and mixing 200 μL fresh media into the well, taking out the media, and putting back 500 μL (movie set 1), or 2) taking out the media, adding back 350 μL and adding 150 μL of fresh media (movie sets 2 and 3). Both methods simulate a media refreshing rate of 0.693/day (equivalent to media half-life = 1 day). For movie sets 2 and 3, an additional centrifuge at around 2000 xg is applied for the old media to remove floating cells. This likely dampened the oscillation in them compared to movie set 1, as iaaH proteins have been shown to work in cell lysate (data not shown).

### METHOD DETAILS

#### Gene constructs

All constructs created in this work were assembled using standard restriction enzyme-based cloning and/or Gibson cloning (Gibson *et al*., 2009). mAID and osTIR1 coding sequences were amplified from addgene #72827 and #72834 (Natsume *et al*., 2016). iCasp9 coding sequence was amplified from addgene #15567 (Straathof *et al*., 2005). PIN2, mGFP, and iaaM coding sequences (CDS) were codon optimized for expressing in mice and synthesized as dsDNA at Integrated DNA Technology together with all oligos for cloning. Coding sequences for screening indole-3-hydrolases were synthesized as cloning plasmid at Twist Bioscience. All constructs were cloned into the piggyBac plasmids (System Biosciences Inc.) driven by a synthetic version of human EF1A promoter.

#### Cell line engineering

All cell lines used in this paper contained stable integrations of transgenes, and were typically clonal populations. To create each stable cell line, the following steps were followed: 1) Cells were first transfected with 800-1000 ng of plasmid DNA using Lipofectamine 2000 or Lipofectamine LTX according to manufacturer’s instructions. 2) 24-48 hours later, cells were transferred to selection media containing 10~50 ug/ml Blasticidin as appropriate for 1-2 weeks. 3) Single clones were isolated through the technique of limiting dilution. For piggyBac constructs, the initial transfection consisted of the target plasmid along with the construct expressing the piggyBac transposase in a 1:4 mass ratio.

#### Flow cytometry

Cells are washed with DPBS (GIBCO) first and then trypsinized with 0.25% trypsin-EDTA (GIBCO) for 1 minute at room temperature. The mixture was then neutralized by culture media and cells were resuspended in Hank’s Balanced Salt Solution (GIBCO) with 2.5 mg/ml Bovine Serum Albumin (BSA). Cells were then filtered through a 40 μm cell strainer (Falcon) and analyzed by flow cytometry (MACSQuant VYB, Miltenyi or CytoFLEX, Beckman Coulter). We used EasyFlow, a Matlab-based software package developed by Yaron Antebi (https://github.com/AntebiLab/easyflow), to process flow cytometry data. All fluorescence data were acquired as the median value of the gated population. For counting cells, 1000 CountBright beads (Life Technologies) were spiked into the sample before filtering, gated out by their fluorescence in analysis and used to estimate cell number in each sample.

#### Conditioned media

The process is described in Figure 2C. Cells were first seeded at about 20% confluence with fresh media for conditioning, and cultured for 2 days. The supernatant was collected as “conditioned media”, and further filtered with 0.22 micron filter or centrifuged at 300 g for 3 minutes to remove any remaining cells. The conditioned media was then combined with fresh culture media at 1:1 ratio, and applied to receiving cells.

#### Cell imaging

For imaging experiments, cells were seeded at 24 or 96-well TC-treated plates with imaging grade plastic bottoms (ibidi inc.), as described above.

Snapshots were acquired on the EVOS imaging system (ThermoFisher) with the EVOS AMG 4x objective (0.13 NA), or a 10x olympus objective (0.30 NA), with the system’s default auto-scanning function.

Time-lapse images were acquired on an inverted Olympus IX81 fluorescence microscope with Zero Drift Control (ZDC), an ASI 2000XY automated stage, iKon-M CCD camera (Andor, Belfast, NIR), and a 20x olympus objective (0.70 NA). Fluorophores were excited with an X-Cite XLED1 light source (Lumen Dynamics). Cells were kept in a custom-made environmental chamber enclosing the microscope, with humidified 5% CO_2_ flow at 37°C. Microscope and image acquisition were controlled by Metamorph software (Molecular Devices). For results in Figure 5A–5C, 12, 25 and 36 positions (about 665um x 665um each) were imaged in each sample, for movie set 1, 2 and 3 respectively. For movie set 1, 12 imaging positions are distributed on a 3×4 grid with 3.5mm intervals. For movie set 2 and 3, the positions are distributed in a 5×5, or 6×6 grid with 600um intervals, and later stitched using Fiji’s stitching plugin (Preibisch, Saalfeld and Tomancak, 2009). Image of each position is acquired every hour for movie set 1, and every four hours for movie set 2 and 3. Figure S2C and movie S1 were imaged with the same method as movie set 1 in Figure 5.

For analysis in Figure 2E, the images were background-subtracted, stitched together and masked by the constitutive mTagBFP2 fluorescent in blue channel (not shown) and quantified by summing up intensities of pixels that passed the mask. Error bars are standard deviations of four images at the same distance.

#### Long-range gradient setup

Silicone-based inserts (ibidi inc. #80269) were first attached to the bottom of TC-treated 6cm dishes (Thermo Scientific). Sender-receiver cells were seeded inside the inserts and allowed to settle down for 2 hours. The inserts are removed and the whole dish is washed with PBS twice to remove non-attached Sender-Receiver cells. Receiver cells were then seeded in the dish at approximately 20% confluence, and allowed to settle down for another 6 hours. To prepare agarose infused media, 2% low melting point agarose (EMD) was melted in alpha-MEM at 95°C for 10 minutes, and temperature was cooled to 42°C, before IAM and other ingredients of complete media (described above) were mixed in. Agarose infused media was applied to dishes with original media removed and allowed to solidify at room temperature for 20 minutes, before moved into the incubator.

#### DNA sequencing of the isolates

Isolates were passaged for at least twice after the long term cultures, before harvested for genomic DNA extraction (Qiagen #69504, with on column RNA digestion). The extracted DNA were then used to amplified the region from the EF1a promoter to the polyA sequence (TaKaRa Bio #R010B) (position 824-8949 on the plasmid map), including all the open reading frames of all the circuit components. Sequencing library were prepped and indexed with Illumina Nextera XT kit (Illuminia #FC-131-1024) and sequenced on Illumina MiSeq, with 300nt pair-ended setup.

#### Bulk RNA sequencing of isolates

Cells were cultured for 2 days at under 80% confluence before harvested for bulk RNA extrusion (Qiagen #74104). The library preparation and sequencing was performed at Millard and Muriel Jacobs Genetics and Genomics Laboratory at Caltech as service. Specifically, RNA integrity was assessed using Bioanalyzer (Agilent Technologies #5067-1513) and mRNA was isolated using NEBNext Poly(A) mRNA Magnetic Isolation Module (NEB #E7490). RNA-seq libraries were constructed using NEBNext Ultra II RNA Library Prep Kit for Illumina (NEB #E7770) following manufacturer’s instructions. Libraries were quantified with Qubit dsDNA HS Kit (ThermoFisher Scientific #Q32854) and the size distribution was confirmed with Bioanalyzer (Agilent Technologies #5067-4626). Libraries were sequenced on Illumina HiSeq2500 in single read mode with the read length of 50nt to the depth of 15 million reads per sample following manufacturer’s instructions. Base calls were performed with RTA 1.13.48.0 followed by conversion to FASTQ with bcl2fastq 1.8.4.

#### Estimation of auxin diffusion coefficient

To estimate the auxin diffusion coefficient, we modeled the system as a fixed concentration source, with the standard result for one-dimensional diffusion:

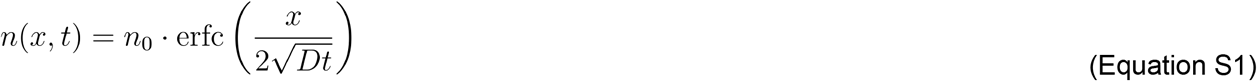

In which, *n* is the concentration as a function of time (*t*) and space (*x*), *n*_0_ is the fixed concentration source, erfc the complementary error function and D is the diffusion coefficient. The diffusion coefficient of auxin has been previously measured to be 3-58 × 10^6^*cm*^2^/*s* (Robinson, Anderson and Lin, 1990). Therefore after 48 hours, the diffusion will expand to approximately 3 to 4 centimeters (Figure S1D). Compared to this ideal fixed concentration source model, the *in vitro* experimental gradient (Figures 2E) is expected to exhibit delays in both auxin production and auxin sensing. Nevertheless, the experimentally observed gradient length scale was similar to the theoretical model prediction (tens of millimeters).

#### Fitting for the auxin and population responsive curves

To fit the mCherry’s response curve to auxin and population (Figure 1C, 3A and 3B; Figure S2A and S2B), we assumed the log of fluorescence (*F*) follows a inverted Michaelis-Menten’s equation:

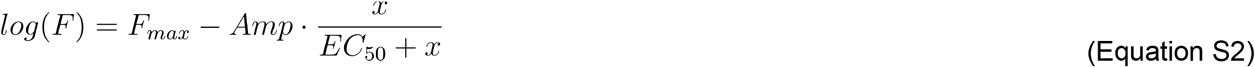

Here, *F_max_* represents the basal level of fluorescence in log; *Amp* represents the max amplitude the population or auxin (represented by *x*) could reduce the fluorescent; and *EC*_50_ represents the concentration, or population number when the reduction is half of the *Amp*.

In Figure S2B, to avoid the problem that the above function (Equation S2) does not extrapolate well when fitted with flat lines (controls and samples treated with auxin), we used *x* = 103 and *x* = 3 · 105 for the extreme values. More specifically, the “ideal dynamic range” was defined as 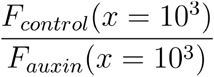, the “loss to self-sensing” was defined as 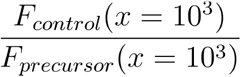, the “loss to saturation” was defined as 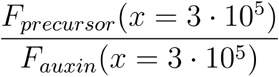, and the “actual dynamic range” was defined as 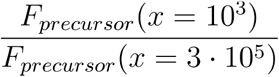.

#### Model of the paradoxical population control circuit

Here we describe a dynamical model of the paradoxical population control circuit. The model is based on the following biochemical reactions, interactions, and assumptions:

- Auxin, denoted *A*, is synthesized from its precursor through an iaaH-catalyzed hydrolysis reaction at a constant synthesis rate per cell *λ_A_*, eliminated at a rate *δ_A_*, which is dominated by dilution due to media changes. Auxin diffuses rapidly in and out of the cell compared to the timescales of cell population dynamics, and its concentration is therefore assumed to be at quasi-steady state (Equation 1).
- iCasp9, denoted *C*, and BlastR, denoted *R*, are produced at rates *λ_C_* and *λ_R_*, respectively.
- Auxin binds reversibly to osTIR1 to form an auxin-osTIR1 complex, which ubiquitylates and degrades iCasp9 and BlastR via their attached AID domains. These reactions are described using classical enzyme kinetics with the auxin-osTIR1 complex as the activating enzyme, described as a constant rate *v_ub_*. In addition to auxin-induced active degradation, iCasp9 and BlastR are also eliminated at rate *δ* due to dilution (Equations S3 and S4).
- The concentrations of iaaH and osTIR1 are assumed to be constant. Auxin precursor is assumed to remain at excess, saturating concentration.
- osTIR1 is assumed to be present at excess concentration compared to the iCasp9-auxin-osTIR1 and BlastR-auxin-osTIR1 complexes, and therefore potential competition between iCasp9 and BlastR for osTIR1 can be neglected.

With these assumptions, we can describe the dynamics of iCasp9 and BlastR with the following differential equations:

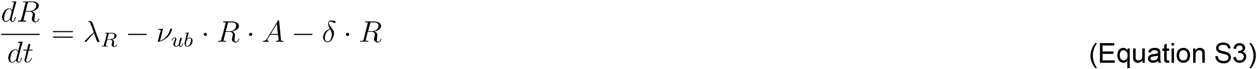

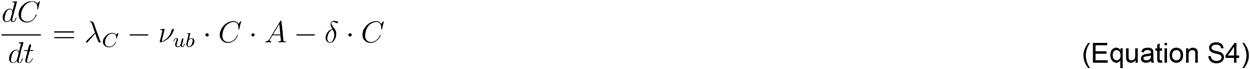

The model represents blasticidin and AP1903 interactions as follows:

- Extracellular Blasticidin, denoted *B*, diffuses into the cell, where it is denoted *B_int_*, and undergoes subsequent enzymatic inactivation by BlastR, with a threshold concentration of *K_B_*, and a Hill coefficient *n*_1_:

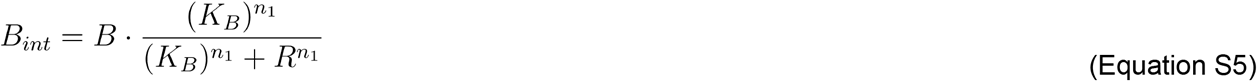
- AP1903 forms an active caspase complex, [*I*: *C*_*n*2_] with iCasp9, with a threshold concentration of 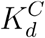 and a Hill coefficient *n*_2_:

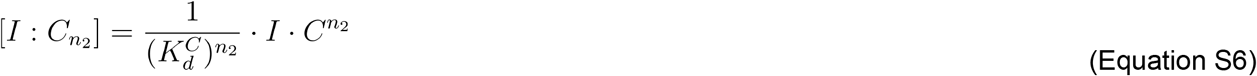

In these two cases, we allow the more general Hill kinetics to account for potential intermediate reaction mechanisms that could influence the effective cooperativity in the final expressions. Additionally, we assumed that both inactivation of Blasticidin and iCasp9 binding to AP1903 are rapid and have reached steady state.

As described in the main text the overall population dynamics can be described using a generalized logistic function, with the growth rate represented as a linear combination of blasticidin-dependent and iCasp9-dependent terms (Equation 2). The Blasticidin-dependent growth rate, *F_G_*, is a sum of two terms. The first describes attenuation of the maximum natural cell proliferation rate, *α*, with increasing blasticidin, while the second represents an increase in the cell death rate, *β*, with increased blasticidin. These terms are associated with half-maximal blasticidin concentrations of *K_g_* and *K_d_* respectively:

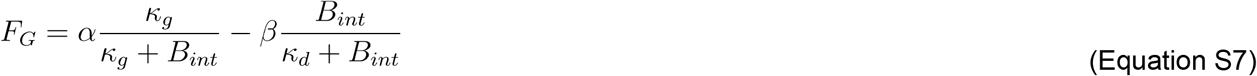

For simplicity, we assume *κ_g_* = *κ_d_* = *κ* Thus, Equation S7 can be reduced to the following form:

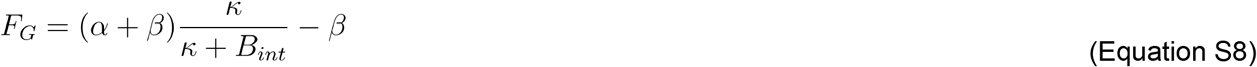

We similarly describe the iCasp9-dependent cell death rate, *F_D_*, with a Hill function dependence on the concentration of the AP1903-iCasp9 (*I: C*_*n*_2__) complex:

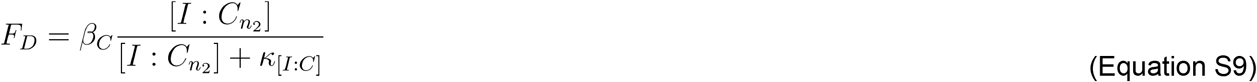

Adding Equations S8 and S9 together and substituting the corresponding terms from Equations S5 and S6, generates the complete form of the growth rate function *R*(*B, I, A*) in Equation 2:

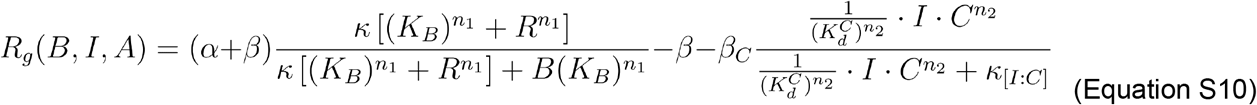

To simplify this description, we assumed a time-scale separation between the faster auxin-population dynamics and the slower intracellular reactions involving *R* and *C*. Using singular perturbation theory (Del Vecchio and Murray, 2014), the system can then be approximated by a simpler system that retains only the slower dynamics (Equation 1 and 2), while the faster dynamics (Equation S3 and S4) are considered to be at equilibrium: 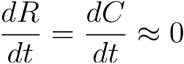. With this approximation, we can write *R* and *C* in terms of the auxin concentration, *A*.

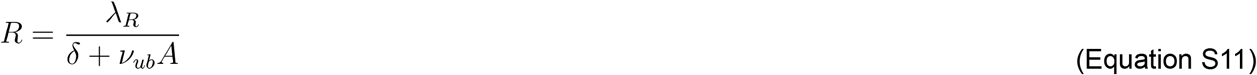

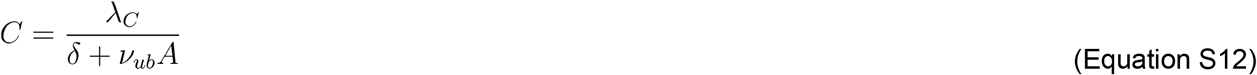

We also defined the following additional parameter combinations for simplicity (see Table S1):

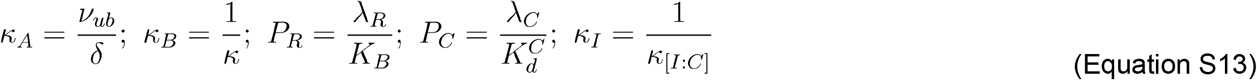

We can then substitute *R* and *C* in Equation S10 to obtain the following:

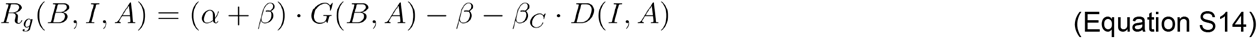

in which:

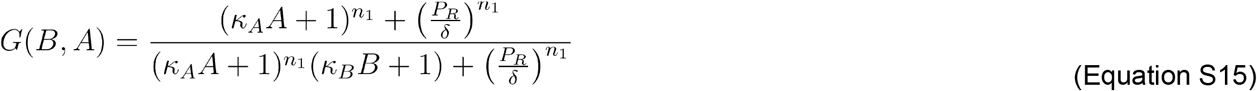

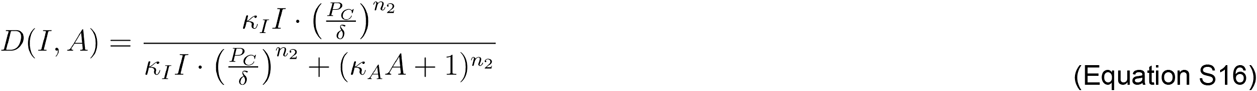

To verify the validity of the approximation, we compared the full model (Equations 1, 2, S3, and S4) to the approximate model, where Equations S3 and S4 are set to 0. We simulated both models using the parameter values in Table S1 (the Paradaux set). The two sets of traces closely followed each other (Figure S3B), indicating that the approximate system accurately reproduced the dynamics of the full model.

#### Representing synergy between iCasp9 and blasticidin control of cell survival

As discussed in the main text, the model above overestimated actual survival rates when both arms of the circuit were simultaneously active (Figures 4D; Figure S3A, dashed purple lines), likely due to synergy between apoptosis and blasticidin-dependent translational inhibition. We therefore added a phenomenological synergistic interaction term: -*β_syn_* · [1 – *G*(*B, A*)] · *D*(*I, A*) to the growth rate expression (*R_g_* in Equation S10):

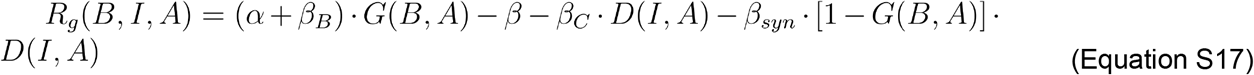

Equation S17 gives the final form of the paradoxical growth curve with the synergistic correction, and is used to improve data fitting (Figures 4D; Figure S3A, dotted purple lines). Together with the fitted parameters, the reduced system (Equations 1 and 2, with *R_g_* defined as Equation S17) was used to run parameter screens and dynamic simulations (Figure 4E–4G, S3E–S3F).

#### Parameter screening and stability analysis

For numerical parameter screening and stability analysis, we computed some terms analytically to make the process faster and more efficient. The stability at equilibrium points of the reduced dynamical system (Equations 1 and 2), is determined by its Jacobian matrix (*J_eq_*, see below) and its eigenvalues.

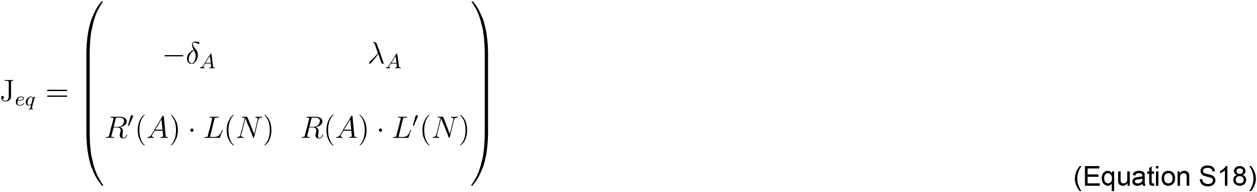

Here, *L*(*N*) = *N*(1 – *N^v^*).

Note that in the operating range (0 < *N* < 1), *L*(*N*) > 0 and *L*’(*N*) > 0; and at any equilibrium points, *dN/dt* = 0. Together with the explicit form of Equation 1, the points above leads to *R*(*A*) = 0 at equilibrium. Therefore, the eigenvalues (*λ*_1_ *and λ*_2_) of at equilibrium are the roots of the quadratic characteristic equation:

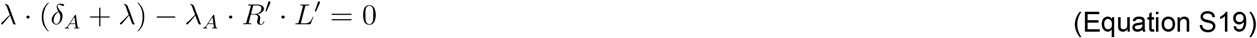

As noted above that *L* > 0, the sign of roots (eigenvalues *λ*_1_ *and λ*_2_) can be solely determined by the sign of *R*’. For *R*’ > 0, at least one of the eigenvalues has a real part greater than zero, making the associated equilibrium point unstable. For If *R*’> 0, we can deduce that the real parts of both eigenvalues are less than zero, making the equilibrium point stable. Based on this analysis, we screened the equilibrium points of the system and the corresponding sign *of R*’ to determine the stability and type of each parameter set (Figure 4E and 4F).

Besides the five major behavior categories described above (Figure 4E, surrounding plots), a small but significant portion (0.68% of total) of conditions appearing at the border between “permissive” and “paradoxical” types, could not be classified into any of the five types. To further investigate these cases we down-sampled the space from 201 x 201 to 21 x 21 conditions and plotted all the six (0.62% or total) unclassified curves, as well the permissive and paradoxical types next to this region (Figure S3G; grey, pink, and light blue respectively). Inspection of these curves revealed a transitional behavior between permissive and paradoxical, with the unusual equilibrium points caused by the nonlinearity of the introduced synergetic term. Note that the range of net growth rate of the system is around −0.09~0.04 defined by *α*, *α* – *β_B_* and *α* – *β_C_*. The net growth rates of these curves around the unusual equilibrium points is significantly lower (−0.01~0.02), indicating those points are unstable. Therefore, those curves’ dynamics, although not mathematically classified, will behave similarly to either permissive or paradoxical types.

#### Delayed blasticidin killing affects oscillations

To simulate the delay effect of blasticidin killing cells (Sato *et al*., 2012), we added a time delay *τ* to the blasticidin related growth function (Equation S15) at time *t*, resulting in a delayed blasticidin growth term, *G*(*B,A*)_*t-τ*_. We then replaced blasticidin related growth term *G*(*B,A*) in Equation S17 with this new term with delay:

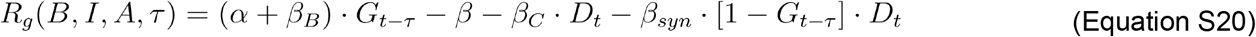

In the above equation, we omitted the arguments of the growth functions for simplicity (e.g. *G*)(*B, A*)_*t*_ is simplified to *G_t_*). Additionally, for simulations in Figure S3F and movie S2 we assumed *G_t-τ_* = *G*_0_; ∀*t < τ*. The results of this delay are the oscillations shown in Figure S3F and movie S2.

### QUANTIFICATION AND STATISTICAL ANALYSIS

#### Confluence estimation

Images, regardless of acquisition conditions, were converted to grey-scale for analysis in cases where pseudo-color was applied by the software automatically (a 2×2 binning is applied if acquired using the EVOS system). For each experiment, around 5 images were used in ilastik (Berg *et al*., 2019) pixel classification mode to train a classifier (decision tree-based), that classifies each pixel as cell or not cell (the trained models are available with the full data set). The classifier was then applied to the entire set of images, and output as probability masks. The masks were then analyzed to determine the fractions of “cell” pixels in the image. This value was then used to estimate confluence.

#### DNA-sequencing analysis

The reads were first trimmed and filtered by Trim Galore! (Babraham Institute) and then aligned with Bowtie2 (Langmead and Salzberg, 2012) to the template sequence (position 824-8949 on the plasmid map). After alignment, we calculated the distribution among four possible nucleotides at each position based on the result from the parental line, as our expected nucleotide distribution for each position (one pseudocount included to avoid significance calling due to low sampling size). We then compared the frequencies of such from other isolates to the expected distribution above to identify any significant deviation (Chi-Square test was used here as a good approximation for multinomial test. p<0.05 with Bonferroni correction). All the significant results are plotted, with their corresponding isolates origin and frequency, in Figure S4A. Among these mutations, we assumed synonymous mutations follow normal distribution to set our expected mutation frequencies for neutral mutations, and determined any non-synonymous mutation that exceeds the 5th percentile (after Bonferroni correction) of this distribution is potentially positively selected (Figure S4A, red dotted line). These mutations are detailed in Table S2.

#### RNA-sequencing analysis

The reads were first trimmed and filtered by Trim Galore! (Babraham Institute) and then counted with kallisto (Bray *et al*., 2016) with the CHO transcriptome (RefSeq GCF_000223135.1) as a reference. Genes that are differentially expressed were determined with DESeq2 (Love, Huber and Anders, 2014) and exported as ranked lists, with *sign*(*log2FoldChange*) · -*log*(*padj*) as weights. The top 30 genes in each of the three comparisons (negative vs control, paradoxical vs control and paradoxical vs negative), if not annotated with a symbol in the RefSeq file, are manually annotated by looking up their gene products in RSCB PDB (RSCB.org) (Basner, 2017). The ranked gene list is then input into the GSEA (Subramanian *et al*., 2005) to perform pathway enrichment analysis, against the curated KEGG pathway annotation (MSigDB: c2.cp.kegg.v7.4.symbols), with mapping from mouse gene symbol to human orthologs.

#### Bootstrapping and significant tests

The curve and significance test in Figure 5C are based on the Kaplan-Meier estimation of the “survival rate”. Between negative and paradoxical feedback, a log-rank test is performed with the null hypothesis that there is no significant difference between the estimated survival rate (Kishore, Goel and Khanna, 2010), using the lifeline python package (Davidson-Pilon, 2021).

For expression ratios (Figure 5F) and survival rates (Figure 5G and 5H), the means and standard deviations were calculated by normalizing treatment replicates to control replicates with bootstrapping. Specifically, for each isolate, we perform replicates for both the treatment (n=3 for 5F-5H) and control (n=3 for 5F, n=2 for 5G-H) group, then normalize the measurements from the treatment groups to the control groups, with all possible combinations (n=9 for 5F, n=6 for 5G-5H). The means and standard deviations of all the normalized measurements were reported as a single data point with bars in the corresponding figures.

Significance tests in Figure 5E–5H are from double-ended Student’s T test, with a null hypothesis that assumes the mean values of the means from each individual isolate are not different.

## Supplementary video and table titles and legends

**Movie S1. Sender-Receiver-PIN2 cells escape after 15 days of continuous culture, related to Figure S2**

Cells were seeded into a 24-well imaging plate with 50 μg/ml of blasticidin solely (“Uncontrolled”) or with 100 μM IAM (“Population circuit on”) or IAA (“Killing constitutive on”). Images of constitutively expressed NLS-citrine were taken with a 20x inverted microscope once per hour.

**Movie S2. Numerical simulation reveals delay bifurcation between damped and limit cycle oscillations, related to Figure 4 and S3**

This movie shows simulated dynamics of the paradoxical feedback model for different values of the delay parameter, *τ.* For each value of *τ,* the simulation shows two trajectories starting from different initial conditions. Note the transition from damped to limit cycle oscillations between 42 and 54 hours. Initial conditions were held fixed at a cell seeding density of 0.1 and auxin concentration of 40 μM and 16 μM for trajectory 1 and trajectory 2, respectively.

**Movie S3. Time-lapse imaging from Figure 5A reveals robustness of the paradoxical population control architecture, related to Figure 5**

10,000 cells were seeded per well into a 24-well imaging plate (Figure 5A), with no control (100μM NAM; left), negative feedback (100 μM NAM and 50 μg/ml blasticidin; middle), or paradoxical feedback (100 μM NAM, 50 μg/ml blasticidin, and 50 nM AP1903; right). Images of constitutively expressed mTagBFP2 were taken with a 20x inverted microscope every 4 hours, in a 6×6 grid, and stitched together. Bar=500micron. This movie is from movie set 3. (Methods).

**Table S1: List of parameters and fitted values, related to Figure 4 and S3**

Fitted values are labeled as (1) or (2), in cases where the major Paradaux line (labeled as 1, Figure 4D), and the one used from confirming the model (labeled as 2, Figure S3C) were fit with different values. Dimensionless values are labeled as “DL” in the “Unit” column.

**Table S2: Details of the mutations detected in DNA sequencing results, related to Figure S4**

Circuit components of isolates from movies set 1 and 2 were amplified and sequenced. For simplicity, uncontrolled growth, negative feedback and paradoxical feedback conditions are noted as unCt, Neg and Para, respectively, in the table. Four out of 18 sequenced isolates exhibited non-synonymous mutations at frequencies that were significantly elevated compared to those of synonymous mutations, consistent with positive selection (Bonferroni corrected, p<0.05, Figure S4A). Three of these (one in a negative feedback cheater isolate, the other two from non-cheating paradoxical isolates) squired mutations in the AID domain of the AID-BlastR fusion protein. Reduced activity of this AID domain would be expected to increase fitness under both paradoxical and negative feedback conditions. The fourth mutation, in a non-cheating paradoxical isolate, was a premature stop codon in osTIR1, near the 5’ end of the circuit transcript, which could potentially eliminate expression of all circuit components from the integrated copy in which it appears. (Because there are multiple genomic integrations of the circuit, any individual mutation could have a limited beneficial effect.)

